# Pan-cancer analysis of patient-specific gene regulatory landscapes identifies recurrent PD-1 pathway dysregulation linked to outcome

**DOI:** 10.1101/2025.10.03.680219

**Authors:** Tatiana Belova, Ping-Han Hsieh, Ladislav Hovan, Daniel Osorio, Marieke L. Kuijjer

## Abstract

Cancer heterogeneity remains a major challenge to the development of broadly effective therapies. Although gene expression-based stratification has advanced our understanding of molecular subtypes, it often overlooks the regulatory mechanisms driving tumor progression and therapeutic resistance. These mechanisms are governed by gene regulatory networks—complex systems of transcription factors and other regulators that control gene expression. To investigate regulatory heterogeneity at the patient level, we inferred individual gene regulatory networks for over 9,000 tumors across 33 cancer types from The Cancer Genome Atlas enabling a comprehensive pancancer analysis of inter-patient regulatory variation. Our analysis uncovered novel regulatory subtypes and revealed recurrent dysregulation of the PD-1 signaling pathway in 23 cancer types. This dysregulation was associated with patient outcomes in 12 cancers and appeared largely independent of established molecular subtypes and immune cell composition, suggesting alternative regulatory mechanisms. We identified both well-characterized (FOS, JUNB, ATF3) and previously unrecognized (RFX7, ELF1, NFYA, NFYC) transcriptional regulators of PD-L1 (*CD274*), highlighting new candidate targets for modulating immune checkpoint activity across tumor types. Our work demonstrates the power of single-sample gene regulatory networks to uncover hidden layers of tumor biology, improve patient stratification, and inform future research in immunotherapy.

## I. INTRODUCTION

Cancer is a highly heterogeneous disease characterized by complex genetic, transcriptional, and regulatory alterations that drive tumor progression and therapeutic resistance. Inter-patient heterogeneity—the variability in molecular features between individuals with the same cancer type—poses a significant challenge to the development of universal therapeutic strategies. Differences in gene expression, mutational landscapes, epigenetic profiles, and tumor microenvironments contribute to variable disease trajectories and treatment responses, highlighting the importance of studying tumor heterogeneity to develop tailored, personalized treatment options and improve therapy selection.

Traditional gene expression-based tumor stratification has provided valuable insights into molecular subtypes, but often fails to capture the regulatory mechanisms that drive oncogenic processes. Expression levels alone do not fully reflect how genes interact in transcriptional networks or how regulatory programs are rewired in cancer. Gene regulatory networks (GRNs) address this limitation by reconstructing transcriptional interactions at a systems level, offering deeper insights into how key transcription factors (TFs) and regulatory elements control oncogenic pathways. For instance, GRN-based approaches have been successfully applied to uncover master regulators of mesenchymal transformation in glioma [1], identify genomic regulatory regions governing proliferative and invasive melanoma states [2], and characterize regulatory differences between intra- and intertumoral regions driving disease progression in colon cancers [3].

Recent advances in single-sample gene regulatory network inference have further expanded the utility of GRNs by enabling the reconstruction of patient-specific transcriptional landscapes. Unlike population-level approaches, single-sample methods capture individual variability in regulatory interactions, allowing for more precise tumor stratification and direct comparisons of regulatory programs across patients. These approaches have been used to characterize regulatory heterogeneity in leiomyosarcoma, revealing novel molecular subtypes [4]; investigate regulatory rewiring associated with survival outcomes in glioblastoma [5]; and explore sex differences in lung adenocarcinoma [6]. Despite these advances, patient-specific GRNs have remained largely unexplored in pan-cancer analyses due to their complexity and largescale nature.

In this study, we performed a large-scale analysis of patient-specific gene regulatory networks across 33 cancer types from The Cancer Genome Atlas (TCGA), comprising over 9,000 primary tumor samples. Our goal was to map regulatory heterogeneity and uncover biologically meaningful tumor subtypes that are not apparent from gene expression data alone. To our knowledge, this is the first comprehensive effort to characterize genomewide regulatory differences at this scale. By inferring regulatory networks for individual patients, we identified patterns of pathway-level dysregulation across tumor types and identified novel, network-based molecular subtypes and regulatory biomarkers (Figure 1).

**Figure 1:**
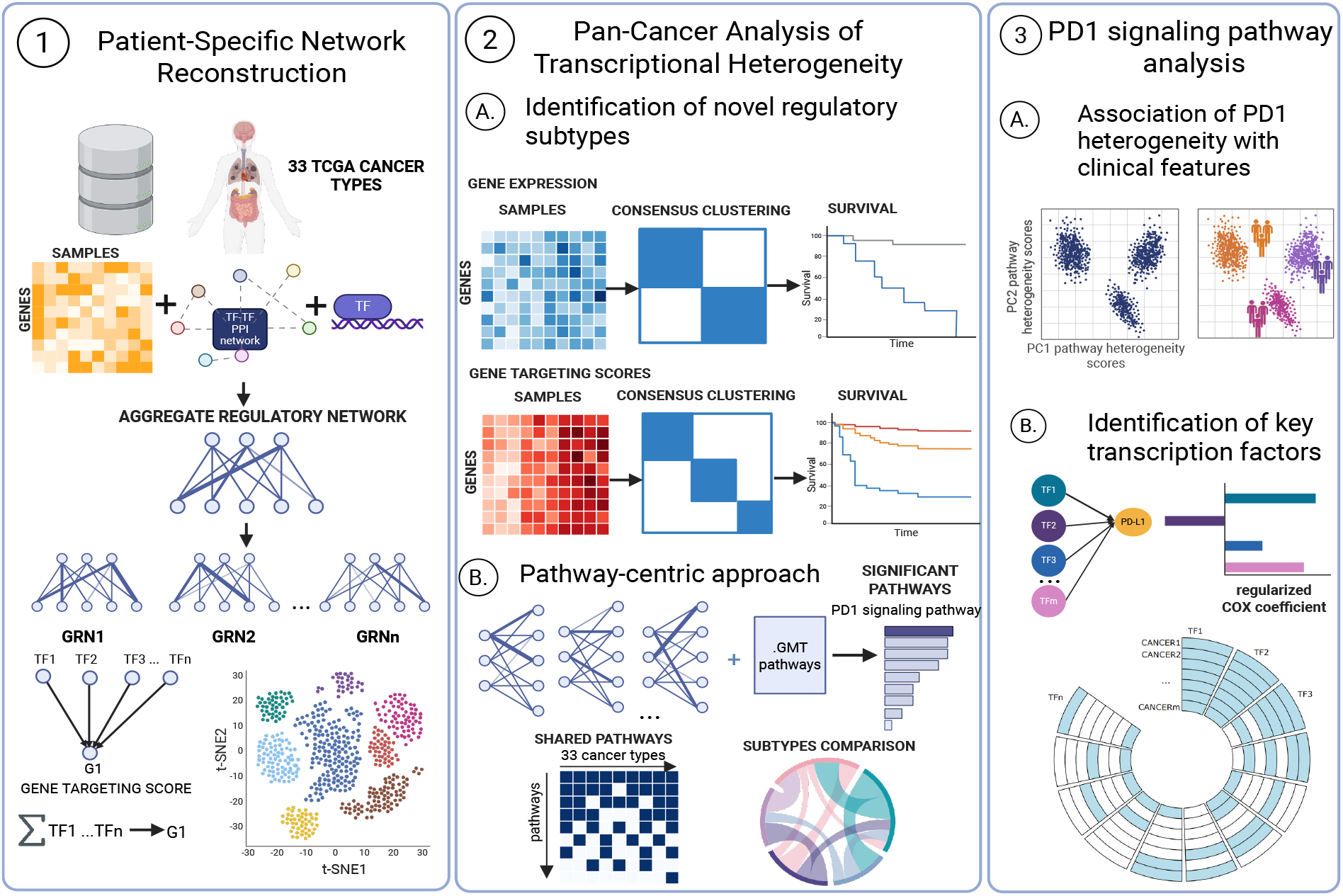
Overview of the pan-cancer analysis of patient-specific gene regulatory networks. Patient-specific gene regulatory networks (GRNs) were reconstructed for over 9,000 primary tumor samples across 33 TCGA cancer types. These GRNs enabled a pathway-centric analysis of transcriptional heterogeneity, leading to the identification of novel molecular subtypes via consensus clustering of gene expression and network-derived gene targeting scores. Downstream analyses uncovered clinically relevant pathways, such as PD-1 signaling, and prioritized key transcription factors associated with regulation of PD-L1 *CD274* and clinical outcome.

The PD-1 signaling pathway—in which PD-L1, encoded by *CD274*, is a key activating ligand—emerged as the most significant finding, exhibiting differential regulation across 23 of the 33 cancer types studied. Its dysregulation was associated with patient prognosis in twelve cancers. This correlated with immune infiltration in only five of them, and appeared largely independent of known molecular subtypes, suggesting that an independent mechanism may be driving this heterogeneity. We further examined the regulatory interactions between the transcription factors controlling *CD274* expression and identified several with potential prognostic relevance. While some of these, including FOS, JUNB, and ATF3, are well-known regulators of *CD274* and tumor progression, our analysis also identified novel associations of *CD274* with RFX7, ELF1, NFYA, and NFYC, uncovering previously unrecognized co-regulatory interactions. Although many of these transcription factors displayed cancer-type specificity, a subset was consistently observed across multiple tumor types, suggesting their potential as regulatory biomarkers and therapeutic targets for immune checkpoint modulation.

## MATERIALS AND METHODS

### Gene expression data pre-processing

We downloaded expression data for all the primary TCGA cases using the “TCGAbiolinks” package in R [7]. We performed group-aware normalization of the expression dataset using “SNAIL” (Smooth quantile Normalization Adaptation for the Inference of coexpression Links) package in Python [8]. SNAIL was specifically designed to improve the accuracy of coexpression measurements, which are leveraged in network reconstruction.

Batch information for TCGA samples was obtained from Molania *et al*. [9]. The presence of batch effect in each cancer was carefully examined with the “PCA-plus” approach using the “MBatch” package in R [10]. This package calculates the distance similarity coefficient (DSC), a metric that quantifies batch effects by comparing the similarity of samples within batches to that across batches. Higher DSC values (above 0.5) indicate stronger batch-specific clustering and, thus, more pronounced batch effects. DSC values were calculated for each cancer type, and cancer types with DSC values above 0.6 were further examined for batch effects (Supplementary Figure S1). Batch effect was identified and corrected using the “sva” package in R in several cancer types, including colon adenocarcinoma, lymphoid neoplasm diffuse large B-cell lymphoma, lung adenocarcinoma, prostate adenocarcinoma, rectum adenocarcinoma and uterine corpus endometrial carcinoma. Batch correction was not applied to esophageal carcinoma, because the observed batch-related separation coincided with the biological distinction between adenocarcinoma and squamous cell carcinoma subtypes.

### Inference of individual patient-specific gene regulatory networks

We used the SPONGE (Simple Prior Omics Network GEnerator) tool to generate a “prior” regulatory network of putative TF-gene interactions and a protein-protein interaction (PPI) network of putative PPIs between transcription factors [11]. SPONGE integrates information from the JASPAR database of transcription factor binding motifs and the STRING database of PPIs to generate these prior networks [12, 13]. The resulting prior networks were filtered by intersecting them with gene expression data to retain only genes and transcription factors with available expression data.

These prior networks were then used as input in PANDA (Passing Attributes between Networks for Data Assimilation) method, using the implementation with GPU acceleration in the netZooPy Python package (https://github.com/netZoo/netZooPy) [14]. PANDA uses a message-passing algorithm to integrate the prior regulatory network, transcription factor PPI data, and gene expression profiles to construct a consensus, or “aggregate,” gene regulatory network. The PANDA output is a complete bipartite graph where each edge represents the inferred regulatory relationship between a transcription factor and a target gene, weighted by the likelihood of the inferred interaction. The PANDA “aggregate” gene regulatory network was estimated based on a total of 9,778 primary tumor samples, for 19,034 genes, and 671 TFs.

To derive individual, sample-specific regulatory networks from this aggregate model, we applied LIONESS (Linear Interpolation to Obtain Network Estimates for Single Samples) [15]. LIONESS iteratively removes each sample from the dataset and recalculates the PANDA network, comparing it to the full aggregate network. LIONESS then uses a linear equation to estimate a unique regulatory network for each individual sample, under the assumption that each edge weight in the aggregate network is a linear combination of its values across all samples.

### t-SNE visualization of gene expression and gene regulatory network clustering

To explore patient similarities and potential clustering patterns in gene expression and gene regulatory networks, we applied dimensionality reduction with t-Distributed Stochastic Neighbor Embedding (t-SNE) [16]. To do so, we first performed dimensionality reduction with principal component analysis, extracting information from the first fifty principal components. These components were then used as input for t-SNE dimensionality reduction, with a perplexity set to 20. We performed t-SNE both on gene expression and on the matrix of gene targeting scores calculated for each gene using the same perplexity value. Gene targeting scores (gene indegree) aggregate all edge weights directed towards a gene, reflecting the extent of regulation the gene receives from the complete set of transcription factors present in the network [17]. These scores have previously been used to identify gene regulatory differences in several studies (e.g., [4, 5]).

### Identification of molecular subtypes based on gene regulatory networks and expression profiles

Molecular subtypes were determined for each cancer type using consensus clustering, implemented in the R package “cola” [18]. For expression and gene indegree profiles, we selected features based on the ATC (ability to correlate to other rows) method which selects features based on the global correlation structure. Then, K-means clustering was applied as the partitioning method, with the number of clusters *K* ranging from 2 to 6. The data was *Z*-score normalized, and consensus clustering involved 1,000 iterations, sampling 80% of the samples at each iteration. The optimal number of subgroups for each cancer type was determined using the “suggest best k” function in the “cola” package, which selects the best *K* based on several clustering quality metrics, including the mean silhouette score, the proportion of ambiguous clustering (PAC) score, concordance, and the Jaccard index.

We compared the clustering results derived from gene regulatory networks and expression profiles using Sankey plots to visualize the relationships between clusters. For cancer types with multiple clustering partitions, the partition with the highest number of clusters (maximum *K*) was selected for visualization.

### LIMMA analysis and preranked gene set enrichment analysis

To identify genes differentially regulated among the subtypes identified based on gene regulatory networks, we employed a Bayesian analysis using LIMMA [19] to compare gene targeting scores while adjusting for patient age at diagnosis and sex. We then performed a preranked gene set enrichment analysis [20] using *t*-statistics obtained from the LIMMA analysis, to identify Reactome gene sets [21] enriched for differentially targeted genes. Gene sets meeting the criteria of a false discovery rate (FDR) below 0.05 and an absolute GSEA enrichment score (ES) greater than 0.5 were considered significantly differentially targeted.

### Pathway-centric pan-cancer analysis of transcriptional regulatory heterogeneity

To capture inter-patient heterogeneity at the gene regulatory level, we used PORCUPINE, our previously developed computational tool [4]. PORCUPINE is an approach based on Principal Components Analysis that can detect the main pathways responsible for the heterogeneity observed among individuals in a given dataset. PORCUPINE evaluates whether a set of variables—such as genes within a biological pathway—shows coordinated variation in their regulation across individuals. It then computes pathway-based patient heterogeneity scores (hereafter referred to as heterogeneity scores), which reflect each individual’s position along a specific principal component (PC) axis that captures the pattern of heterogeneity in that pathway. We applied PORCUPINE to gene regulatory networks for each cancer type. To identify the most significant pathways, we applied stringent filtering using three different criteria—1. p-values adjusted for multiple testing with the BenjaminiHochberg method [22] less than 0.05; 2. pathways with effect size larger than the 95th percentile; and 3. the variance explained by the first principal component above 25%.

### Association of PD-1 pathway regulatory heterogeneity with clinical features and cell composition

Curated and filtered clinical information, including outcome endpoints, was obtained from Liu, *et al*. (2018) [23]. Additional clinical data, including subtype details and previously defined molecular “omics” subtypes, was retrieved for each cancer type using the “TCGAbiolinks” package in R. The corresponding DOIs for each cancer type, as provided by “TCGAbiolinks”, are listed in Supplementary Table S1. For certain cancer types, the clinical data also included estimated immune and stromal scores, derived from gene expression profiles using the ESTIMATE algorithm [24]. To investigate the association between heterogeneity in PD-1 pathway regulation and clinical features, we employed different statistical methods based on feature types. For numerical features, we calculated the Spearman correlation coefficient, while for categorical features, we performed pairwise group comparisons using the Mann-Whitney test (Wilcoxon rank-sum test) and measured effect sizes with rank-biserial correlation between the feature and the PD-1 pathway-based patient heterogeneity scores. For each comparison, we extracted the corresponding p-values and effect sizes. P-values from all comparisons were adjusted for multiple testing using the FDR correction with the Benjamini-Hochberg method.

To investigate whether dysregulation of the PD-1 signaling pathway was driven by differences in cell composition, or rather independent of it, we estimated the fractions of tumor-infiltrating cells with CIBERSORTx. To do this, gene expression profiles were used in cibersortx apptainer (http://cibersort.stanford.edu/), where the algorithm was executed with the LM22 signature matrix, employing 1000 permutations. We then used Spearman correlation between immune cell infiltration scores from CIBERSORTx and PD-1 pathway-based patient heterogeneity scores obtained with PORCUPINE. Associations with absolute Spearman correlations above 0.4 were considered significant.

### Identification of key *CD274*-associated transcription factors through regularized Cox regression

To identify the most important regulatory interactions associated with clinical endpoints between all 671 transcription factors and the *CD274* gene, we performed a regularized Cox regression analysis on network edge weights to *CD274* in each cancer type. Regularization applies a penalty to the regression coefficients, shrinking less informative predictors toward zero and effectively selecting a subset of predictors by retaining only those with nonzero coefficients in the final model. We employed elastic net regression using the “glmnet” package in R, setting the alpha parameter to 0.3. We incorporated five-fold cross-validation and repeated the model fitting 100 times to ensure robust and reliable results. In each run, the optimal lambda value with the minimum crossvalidation error was selected and used to fit the final model to select the most predictive features. Only features that had coefficients in at least 50 runs were selected for further analysis. We followed the recommendations of Liu et al. [23] and excluded the pheochromocytoma and paraganglioma (PCPG) dataset due to its limited number of events (only six). For most cancer types, overall survival (OS) was used as the disease outcome. However, for BRCA, LGG, PRAD, READ, TGCT, THCA, and THYM, we used progression-free interval (PFI), as recommended in [23].

## RESULTS

### Patient-specific regulatory networks reveal cancer type-specific and pan-cancer regulatory patterns

We reconstructed patient-specific genome-wide regulatory networks by incorporating expression data for 19,034 genes measured across 9,778 primary tumors derived from 33 cancer types from the TCGA dataset. Using the PANDA algorithm for gene regulatory network reconstruction, we integrated gene expression information from these datasets with an initial prior gene regulatory network of 671 transcription factors and their proteinprotein interactions. We then used LIONESS to obtain sample-specific networks.

First, we investigated whether the reconstructed patient-specific genome-wide regulatory networks reflect known heterogeneity among different cancer types. Using *t*-distributed stochastic neighbor embedding (tSNE) visualization of gene targeting scores (a measure of gene regulation, see Methods) from all samples, we observed that samples from the same tumor type tend to cluster together, showing that genes have distinct cancer type-specific targeting patterns (see Figure 2B). Moreover, kidney cancer types—kidney renal papillary cell carcinoma (KIRP) and kidney chromophobe (KICH), were often in proximity on the t-SNE visualization. Gastrointestinal tumors were also co-localized, with stomach adenocarcinoma (STAD) and esophageal carcinoma (ESCA), as well as colon adenocarcinoma (COAD) and rectum adenocarcinoma (READ) located close to each other.

**Figure 2:**
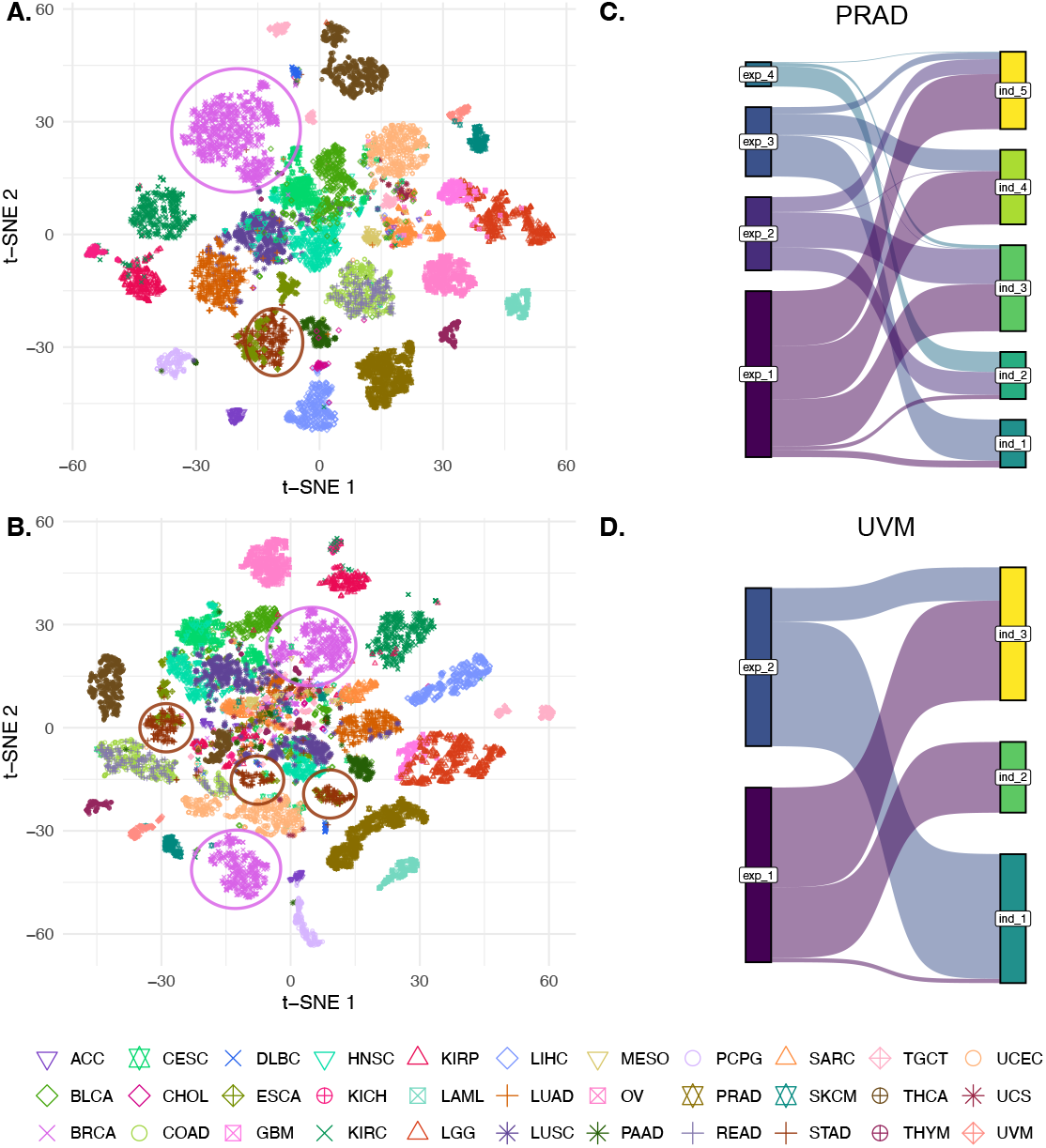
t-distributed Stochastic Neighbor Embedding (t-SNE) visualization of 9,778 primary tumor samples across 33 cancer types, represented by distinct colors and shapes. Panels show clustering based on (A) gene expression and gene targeting scores. The abbreviations for cancer types and their corresponding full names are provided in Supplementary Table S1. Comparisons of “cola”-derived clusters based on gene expression (exp) and network indegree measures (ind) are shown for (C) Prostate Adenocarcinoma (PRAD) and (D) Uveal Melanoma (UVM).

Unlike the visualization observed based solely on expression data (see Figure 2A), several cancer types, such as breast invasive carcinoma (BRCA) and stomach adenocarcinoma (STAD), displayed significant spread and multiple potential clusters, suggesting the presence of regulatory subtypes (shown with circles in Figure 2B). Some cases from different cancer types co-localized at the center of the plot, indicating shared regulatory features that may reflect a more generic, pan-cancer gene regulatory network. This suggests that certain tumors, despite originating from different tissues, may share underlying transcriptional regulatory mechanisms.

### Network-based stratification of cancer provides new molecular subtypes

We next investigated whether tumors from each of the 33 TCGA cancer types can be separated into well-defined subtypes based on their gene regulatory and expression profiles. To this end, we performed a consensus clustering partitioning using *K*-means on each cancer type (see Methods). For various cancer types, several optimal values of *K* were identified (Supplementary Figure S2, S3) and included in the downstream analysis. In general, the clusters derived from the expression data and gene regulatory networks were different from one another (example shown in Figure 2C), all cancer types in Supplementary Figure S4). Some cancer types showed fewer rearrangements, such as uveal melanoma (UVM, Figure 2D).

To evaluate whether patient clusters, identified based on expression and gene regulatory networks, exhibit associations with clinical outcome, we performed univariate Cox regression analyses within each cancer type, adjusting for patient age and sex. Depending on the specific cancer type, outcomes were assessed using either overall survival (OS) or progression-free interval (PFI), as appropriate (see Methods). Significant associations between clusters and clinical outcomes were identified across several cancer types, both for clusters derived from expression and for those derived from gene targeting scores (Supplementary Figure S5).

Notably, for prostate adenocarcinoma (PRAD), we observed significant differences in the network-derived clusters, but not in clusters that were based on expression data (Supplementary Figure S5). To identify regulatory differences between these clusters, we performed a Bayesian statistical analysis (LIMMA). Subsequent gene set enrichment analysis using the Reactome canonical pathway collection from MSigDB [21] revealed pathways involving key processes involved in cancer development and outcomes, such as proliferation, immune signaling, metabolism, and DNA repair (Supplementary Figure S6). The pathway ‘Activated *PKN1* stimulates transcription of AR-regulated genes *KLK2* and *KLK3*’ was identified as one of the top differentially regulated between clusters 1 and 2, which exhibit significant differences in progression-free intervals (PFI), highlighting the known role of androgen receptor (AR) activity in prostate cancer progression and confirming that gene regulatory network-based clustering can capture biologically and clinically meaningful differences that are not evident from expression data alone.

### Network-based pathway-centric analysis highlights common and unique dysregulation in cancer

Encouraged by these findings, we next set out to perform a comprehensive genome-wide analysis of pathway-based regulatory heterogeneity in cancer, aiming to determine whether the observed inter-patient heterogeneity could be further explained by coordinated pathway-level dysregulation. To this end, we used PORCUPINE, a computational tool designed to identify pathways whose genes show coordinated regulatory variability across a population of patient-specific gene regulatory networks [4].

Applying PORCUPINE across all 33 TCGA cancer types with the C2 curated gene sets from the Molecular Signatures Database [21], we uncovered a range of significant pathways displaying differential regulation in each cancer type. Top gene sets were selected based on an FDR *<* 0.05, the variance explained by the first principal component above 25% and effect sizes above the 95th percentile. Jaccard similarity analysis across cancer types revealed high pathway-level similarity among biologically related cancers, confirming our findings from the t-SNE visualization on gene targeting scores (Supplementary Figure S7).

To determine patterns of pathway dysregulation, we next grouped the significant pathways into subcategories according to their cellular function (see Figure 3A). Pathways with genes involved in immune system, cell cycle, signal transduction and disease were differentially regulated in most cancer types—all cancer types have at least one of these, and 85% have two or all three categories affected. A number of categories and pathways were identified as heterogeneous in a more specific set of malignancies. For example, neuronal system pathways were mostly identified in glioblastoma multiforme (GBM) and lower-grade glioma (LGG), digestion and absorption pathway were specifically captured in cholangiocarcinoma (CHOL), while autophagy was specifically identified in chromophobe renal cell carcinoma (KICH). These pathways likely reflect tissue-specificity. Pathways that were shared among at least six cancer types are visualized in Figure 3B. Of note, with this definition of recurrence, the “signal transduction” category did not include recurrent pathways, indicating that dysregulation of signal transduction pathways is often cancer type-specific. The top three recurrent pathways include PD-1 signaling, the “endosomal vacuolar pathway,” and FC gamma receptor protein (FCGR) activation, differentially regulated in 23, 16, and 14 cancer types, respectively.

**Figure 3:**
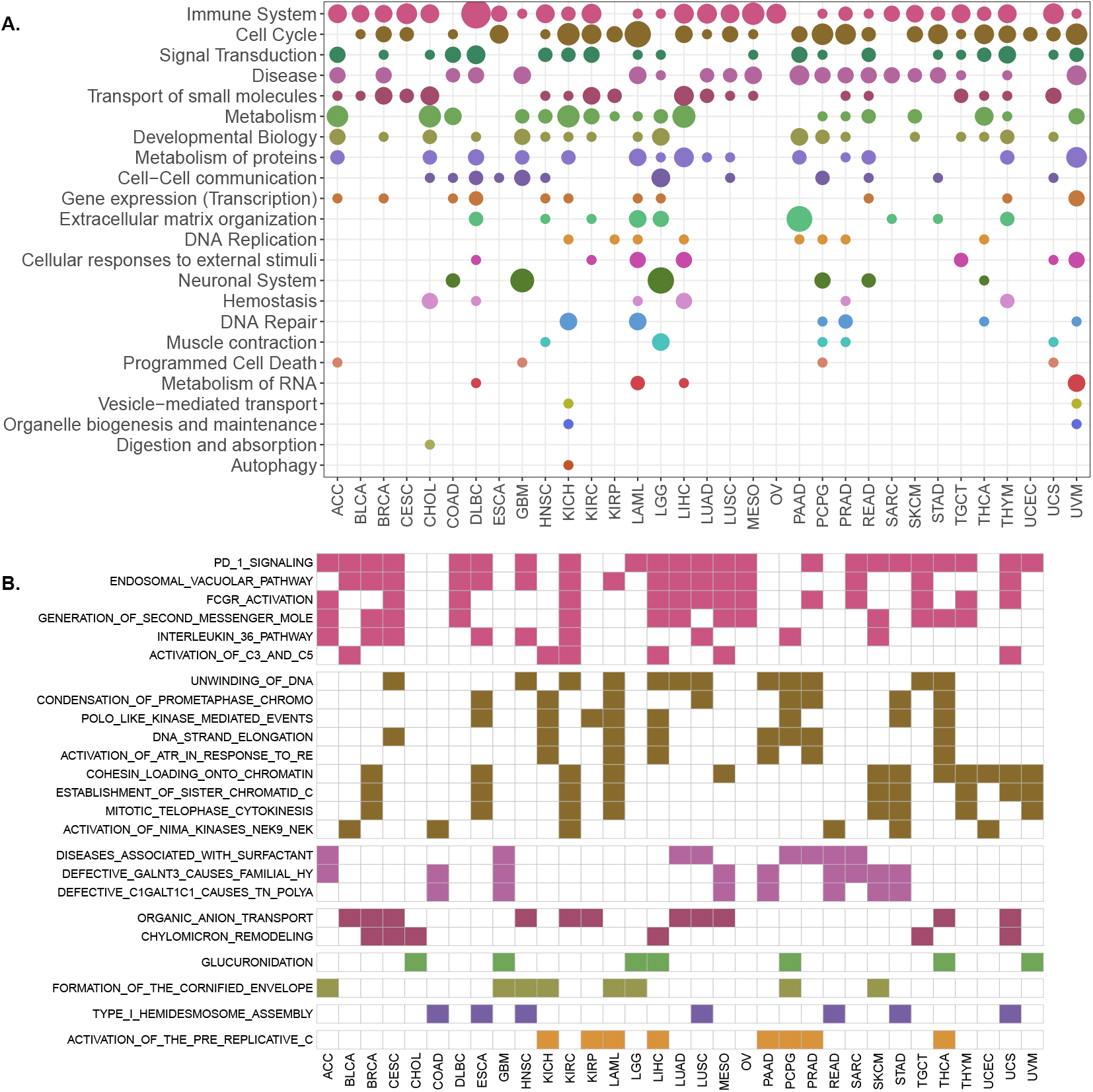
A. Pan-cancer overview of significantly differentially regulated pathway categories, as identified by PORCUPINE (FDR *<* 0.05, variance explained by the first principal component above 25% and effect sizes above the 95th percentile). The circle size represents the percentage of heterogeneous pathways among all the C2 pathways within that cellular category. B. Pathways that exhibit regulatory heterogeneity (colored boxes) in at least six cancer types. Colors indicate a significant association, and correspond to the categories in A.

### Regulation of PD-1 signaling is associated with clinical outcomes across multiple cancer types

As PD-1 signaling was identified as the top differentially regulated pathway across 23 out of 33 TCGA cancer types (see Figure 3B), we next investigated whether this regulatory heterogeneity was associated with clinical outcome. We used the PD-1 pathway heterogeneity scores derived from the first two principal components in a univariate Cox regression model to predict outcome (PFI or OS, depending on the cancer type, see Methods). Significant associations with outcome were observed in 12/32 cancer types (Table 1); note that one cancer type was excluded from the survival analysis, see Methods).

**Table 1.** Summary of PD-1 signaling pathway analysis. A checkmark in the *Heterogeneity* column indicates cancer types in which PORCUPINE analysis identified the pathway as differentially regulated, with this regulatory heterogeneity associated to clinical outcomes (*Clinical outcome* column). For these cancer types, further analyses examined associations between heterogeneity and PD-L1 (*CD274*) gene expression (Spearman correlation; \*\*\**p* − *value <* 0.001), clinical features, previously defined “omics” clusters, and immune cell infiltration.

| Cancer | Heterogeneity | Clinical outcome | PD-L1 expression | Clinical features | Omics clusters | Immune Infiltration (CIBERSORTx) |
| --- | --- | --- | --- | --- | --- | --- |
| ACC | ✓ | ✓ | R = 0.78*** | ✓ | ✓ | ✓ |
| BLCA | ✓ | - |  |  |  |  |
| BRCA | ✓ | ✓ | R = 0.28*** | - | - | - |
| CESC | ✓ | ✓ | R = 0.45*** | - | - | ✓ |
| CHOL | - |  |  |  |  |  |
| COAD | - |  |  |  |  |  |
| DLBC | ✓ | - |  |  |  |  |
| ESCA | ✓ | - |  |  |  |  |
| GBM | - |  |  |  |  |  |
| HNSC | ✓ | - |  |  |  |  |
| KICH | - |  |  |  |  |  |
| KIRC | ✓ | ✓ | R = 0.15*** | - | ✓ | ✓ |
| KIRP | - |  |  |  |  |  |
| LAML | - |  |  |  |  |  |
| LGG | ✓ | ✓ | R = 0.56*** | ✓ | ✓ | - |
| LIHC | ✓ | ✓ | R = 0.23*** | - | - | - |
| LUAD | ✓ | ✓ | R = 0.32*** | - | - | - |
| LUSC | ✓ | - |  |  |  |  |
| MESO | ✓ | - |  |  |  |  |
| OV | ✓ | ✓ | R = 0.25*** | - | - | - |
| PAAD | - |  |  |  |  |  |
| PCPG | - |  |  |  |  |  |
| PRAD | ✓ | ✓ | R = 0.37*** | - | - |  |
| READ | - |  |  |  |  |  |
| SARC | ✓ | ✓ | R = 0.31*** | ✓ | ✓ | ✓ |
| SKCM | ✓ | ✓ | R = 0.41*** | - | - | ✓ |
| STAD | ✓ | ✓ | R = 0.2*** | - | - | - |
| TGCT | ✓ | - |  |  |  |  |
| THCA | ✓ | - |  |  |  |  |
| THYM | ✓ | - |  |  |  |  |
| UCEC | - |  |  |  |  |  |
| UCS | ✓ | - |  |  |  |  |
| UVM | ✓ | - |  |  |  |  |

In the tumor microenvironment, PD-1 pathway activation is typically driven by interactions between PDL1 expressed on the tumor or immune cell and PD-1 on infiltrating T cells [25]. Since *CD274* encodes PD-L1, we assessed whether the PD-1 pathway heterogeneity scores identified with PORCUPINE correlated with PD-L1 (*CD274*) expression levels. This provides complementary information, as our network models capture inferred regulatory relationships rather than direct gene–gene expression correlations. For this, we focused on the twelve cancer types in which PD-1 regulatory heterogeneity was prognostic. We observed significant correlations (Spearman *R >* 0.4, *p* − *value <* 0.001) between *CD274* expression and patient distribution along the respective principal component axes in 4/13 cancer types—ACC (R = 0.78), CESC (R = 0.45), LGG (R = 0.56), and SKCM (R = 0. 41) (Table 1, Supplementary Figure S8).

Notably, in LGG, higher *CD274* expression was associated with worse outcomes, potentially reflecting immune evasion in the context of this typically immune cold tumor. In contrast, in ACC, CESC, and SKCM, higher *CD274* gene expression levels correlated with more favorable outcomes, potentially reflecting an active adaptive immune response in which PD-L1 is upregulated in response to immune cell infiltration—a pattern often linked to improved prognosis (Supplementary Figure S8).

For the remaining cancer types, we did not observe strong correlations between *CD274* expression and PD1 pathway regulatory heterogeneity, suggesting that the regulatory variation captured by PORCUPINE may reflect broader or more complex immune regulatory programs beyond PD-L1 expression alone.

### Regulatory heterogeneity of the PD-1 pathway is largely independent of known molecular cancer subtypes

Previous TCGA studies provide comprehensively curated clinical and pathological features of tumors, as well as molecular classifications based on large-scale multiomics profiling data. To examine whether the PD-1 pathway regulatory heterogeneity captured by our model extended beyond known tumor stratification, we tested its associations with clinical features and molecular subtypes (see Supplementary Table S1).

We restricted our analysis to the cancer types in which PD-1 pathway regulatory heterogeneity was significantly associated with clinical outcome (see Table 1, Supplementary Figure S8). Among these, we identified significant (*FDR <* 0.01, and an absolute Spearman correlation or rank-biserial correlation above 0.4) associations with clinical or molecular features in 4/12 cancer types, including ACC, KIRC, LGG, and SARC.

In adrenocortical carcinoma (ACC) higher PD-1 pathway heterogeneity scores, indicative of a better prognosis, were associated with the C1B subtype, consistent with previous knowledge that classify C1A as aggressive and C1B as indolent (see in Supplementary Figure S12). In kidney renal clear cell carcinoma, PD-1 pathway heterogeneity scores were associated with previously defined classification by expression. In sarcomas, higher scores were associated with worse overall survival, with the leiomysarcoma subtype, higher tumor purity (R = 0.66), higher cancer DNA fraction (R = 0.68), and lower leukocyte fraction (R = −0.78) (see Supplementary Figures S14 and S15).

Finally, in lower-grade glioma, heterogeneity scores were associated with various subtypes, including IDH 1p19q co-deletion subtype, transcriptome subtype classification, and IDH-specific RNA and methylation classifications, as shown in Supplementary Figures S11 and S13. Notably, higher PC1 heterogeneity scores were associated with worse outcome, specifically progression-free interval (PFI), and correlated with higher stromal (Spearman R = 0.45) and immune scores (Spearman R = 0.59) (see Supplementary Figures S12). Tumors belonging to these higher scores were enriched for IDH wild-type compared to IDH-mutant, corroborating the known fact that IDH-mutant tumors generally display a more indolent behavior.

For the remaining eight cancer types, we did not identify strong associations between PD-1 pathway heterogeneity and clinical or molecular classifications (see Supplementary Figure S9), reinforcing the notion that PD-1 signaling heterogeneity represents an independent layer of tumor variability, likely driven by complex regulatory mechanisms beyond established classifications.

### Dysregulation of PD-1 pathway is associated with immune infiltration in some but not all cancers types

While general immune or stromal scores can provide a rough estimate of immune activity within tumors, they may not capture the full complexity or specificity of the tumor immune microenvironment. To obtain a more refined view of immune cell composition, we therefore applied CIBERSORTx deconvolution to estimate the proportions of immune and immune-related cell types in each sample.

We calculated Spearman correlations between PD-1 pathway heterogeneity scores and the estimated immune cell compositions (Supplementary Figure S10). We identified significant associations (*R >* 0.4) between PD-1 pathway heterogeneity scores and CD8+ T-cell proportions for 5/12 cancer types. In addition, for certain cancer types, we found associations with regulatory T-cells (KIRC) macrophages (M1 in ACC and CESC, M2 in ACC), and activated dendritic cells (ACC).

Notably, among the cancer types where PD-1 pathway heterogeneity was significantly associated with PD-L1 (*CD274*) expression and better clinical outcome—namely ACC, CESC, and SKCM—we also observed strong correlations with CD8+ T-cell infiltration. This pattern suggests that, in these cancers, regulatory heterogeneity in the PD-1 pathway may reflect adaptive immune responses involving cytotoxic T-cell activity. In contrast, in LGG, where *CD274* expression was also associated with clinical outcome but showed no correlation with CD8+ T-cell infiltration, PD-1 pathway variation may arise from tumor-intrinsic regulatory programs rather than direct T cell–mediated immune pressure.

### Key regulatory interactions of *CD274* associated with the clinical outcomes

To gain deeper insights into PD-1 pathway regulatory heterogeneity and its relevance for clinical outcomes, we next focused on the transcriptional regulators of PD-L1, a key immune checkpoint ligand with direct implications for tumor immune evasion and immunotherapy response. We again limited our analyses to cancer types that showed significant associations with clinical endpoints (see Table 1 and Figure 4). Specifically, to identify key transcription factors linked to outcomes in each cancer type, we conducted regularized Cox regression analysis on edge weights from all 671 TFs to the *CD274* gene, selecting those regulatory interactions that demonstrated a robust association with outcome, i.e. in at least 50 out of 100 model runs (for results with a more relaxed threshold, see Supplementary Figure S16).

**Figure 4:**
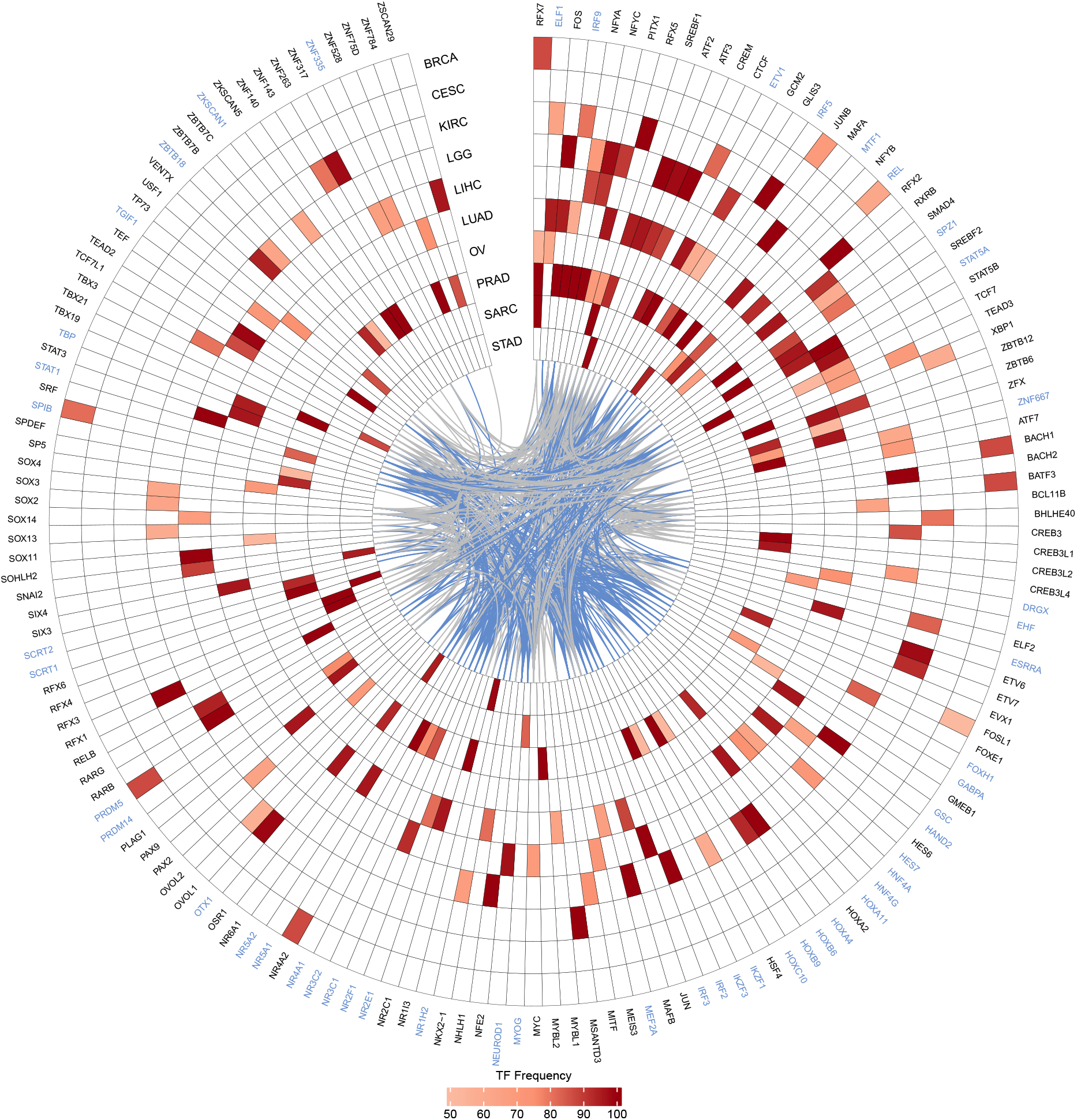
TFs selected through the application of the regularized Cox regression model to edge weights between 671 TFs and the *CD274* gene across various cancer types. TFs with non-zero coefficients were selected and were predictive in at least 50 out of 100 model runs. The color of each cell reflects the frequency with which the TF was selected as predictive. Grey inner connections represent known interactions among TFs based on protein-protein interaction (PPI) data, while blue connections indicate TFs predicted to regulate *CD274* according to the motif prior network. Blue TF names indicate these TFs have prior binding sites in the promoter of *CD274*.

Out of the 13 cancer types analyzed, ten had at least one feature associated with outcome. In total, we identified 160 predictive TF-*CD274* interactions— 51 of these were supported by known TF-binding sites in the *CD274* promoter, which is significantly higher than expected by chance (p-value = 0.04). The cancer types with the highest number of predictive regulatory interactions were LGG, LUAD, and PRAD, with 38, 42, and 51 transcription factors identified, respectively. In contrast, CESC had the fewest features, with only one transcription factor (XBP1) meeting the stringent threshold.

Most of the transcription factors identified were specific to a particular cancer type (see Figure 4). However, we also identified recurrent transcription factors. While some of these, such as ELF1 or IRF9, have known binding sites to the *CD274* promoter, several were not included with a potential binding site in the prior network, likely due to stringent filtering criteria applied during its construction. Nevertheless, various of these are well-documented in the literature as PD-L1 regulators. These include members of the Activator protein-1 (AP1) complex, a transcription factor complex, which is formed by a group of transcription factors comprising four primary subfamilies: the JUN, FOS, MAF, and ATF (activating transcription factor) protein families [26– Several transcription factors, including members of the FOS and JUN families, were identified as significant prognostic markers across multiple cancer types (see Figure 4).

Similarly, several STAT family transcription factors were associated with *CD274* regulation and prognosis, including STAT1, STAT3, STAT5A, and STAT5B across lung adenocarcinoma, liver hepatocellular carcinoma, and sarcomas (Figure 4). Extensive literature highlights the key role of STAT1 in CD274 regulation across multiple cancers, and it has been shown to directly bind the PD-L1 promoter [29– In addition, STAT3 can directly regulate *CD274* by binding to its promoter or indirectly through various signaling pathways, as evidenced in multiple cancer types [29, 35–40]

The most recurrent TFs (selected in at least 3/10 cancer types) included RFX7, ELF1, FOS, IRF9, NFYA, NFYC, PITX1, RFX5 and SREBF1 (Figure 4). Among these, IRF9 and FOS have well-established roles in the transcriptional regulation of PD-L1. As previously mentioned, FOS, as part of the AP-1 complex, contributes to PD-L1 upregulation through MAPK pathway activation, a mechanism frequently observed in various tumor types [27, 41]. Several studies have shown that IRF9 is involved the regulation of PD-L1 expression through both direct transcriptional activation in response to type I IFN signaling and indirect mechanisms involving chromatin organization [42–44]

Overall, this analysis highlights both well-known and previously unrecognized transcription factors, providing new insights into the complex regulatory landscape of PD-1 signaling across different cancer types, with implications for improving patient stratification and pointing to transcriptional regulators that could serve as targets for modulating immune checkpoint activity.

## Discussion

Understanding patient-specific gene regulation is essential for interpreting tumor heterogeneity and its impact on clinical outcomes. By systematically mapping gene regulatory heterogeneity across 33 TCGA cancer types for over 9,000 patients, we provide a comprehensive view of regulatory variation and its relevance for tumor biology and immune regulation. Our findings demonstrate that network-based stratification offers a distinct and biologically meaningful classification of tumors, revealing hidden regulatory patterns not captured by gene expression-based clustering, such as those we identified in prostate cancer.

To further characterize the biological processes underlying this regulatory heterogeneity, we performed a pathway-centric, pan-cancer regulatory network analysis. While some pathways showed marked cancer-type specificity, others were recurrently dysregulated across multiple cancer types. The PD-1 signaling pathway emerged as the most frequently dysregulated, exhibiting significant heterogeneity in 23 out of 33 cancer types. In addition to its high prevalence, this pathway was prognostically relevant in about 40% of all cancer types, underscoring its potential as a biomarker for clinical outcomes and a target for immunotherapy.

Building on these findings, studying the dysregulation of the PD-1 pathway is especially crucial, given that PD-1 and its ligand PD-L1 act as critical negative regulators at the immune checkpoint. Their interaction suppresses cytotoxic T-cell function and promotes tumor immune evasion, highlighting the pathway’s therapeutic importance. Although PD-L1 (*CD274*) expression on tumor cells is widely recognized as a biomarker for immunotherapy response, the underlying regulatory mechanisms that control its expression remain incompletely understood. Gaining a comprehensive understanding of these regulatory mechanisms is therefore beneficial for the immunotherapy strategy.

Since PD-L1 can be expressed by both tumor and immune cells, we performed deconvolution of the RNAseq data into immune cell types to gain a clearer understanding of the immune microenvironment and patterns of immune infiltration. PD-L1 expression on tumor cells typically indicates an intrinsic mechanism of immune evasion, whereas its expression on immune cells is more often linked to the presence of active immune infiltration and an adaptive resistance response. In a subset of cancer types regulation of PD1 signaling was associated with survival (5/12), PD-1 pathway heterogeneity correlated with CD8+ T-cell infiltration, consistent with immune-responsive phenotypes. However, in others, such as lower-grade glioma, regulatory variation occurred without corresponding T-cell infiltration, potentially reflecting tumor-intrinsic mechanisms of immune evasion.

A key observation in our study was that PD-1 pathway–based tumor stratification did not simply reflect existing molecular or clinical subtypes. In most cancer types, regulatory heterogeneity captured by our models was independent of established clinical classifications and “omics” subtypes, and was not explained by PD-L1 expression levels alone. Thus, these could potentially be novel, independent biomarkers of outcome. However, in certain cancer types such as lower-grade glioma, PD-1 pathway–based stratification aligned with known molecular subtypes, such as the IDH-mutant subtype. This suggests that in some contexts, regulatory heterogeneity in immune signaling may coincide with broader genomic alterations, whereas in others, it represents a distinct and clinically relevant axis of heterogeneity. Experimental validation will be essential to translate these networkbased findings into actionable biomarkers for tumor stratification and therapy guidance.

To further dissect the regulation of PD-L1-encoding gene *CD274*, we focused on its upstream transcriptional control and identified regulators for which their interactions with *CD274* associated with clinical outcome. Various of the identified TFs are well-known for their roles in regulating cell proliferation, metabolism, and immune evasion. While several of the identified TFs are direct regulators of *CD274*, we also uncovered novel associations without direct evidence of promoter binding. These transcription factors may modulate *CD274* expression indirectly, through co-regulation, chromatin remodeling, or enhancer-promoter binding. Notably, our approach recovered several well-documented regulators absent from the prior binding network, underscoring the ability of GRN-based inference to identify context-specific or non-canonical regulators.

Although most of the identified TFs were cancer type-specific, a subset was recurrent across multiple tumor types. These recurrent TFs may represent core regulators of *CD274* regulation and tumor immune evasion, making them particularly interesting candidates for further investigation. Their mechanisms of action and their potential for pan-cancer therapeutic targeting merit further investigation. These TFs could, for example, be targeted directly, or modulated indirectly through upstream signaling pathways (e.g., via small molecule inhibitors), offering new approaches that synergize with checkpoint inhibition. In fact, several are already being explored in immuno-oncology, such as blocking activation of STAT3 [45], inhibiting nuclear translocation in NF*κ*B [46], disrupting protein stabilization in HIF-1*α* [47], suppressing transcriptional activity in MYC through bromodomain inhibition [48], and modulating AP-1 activity via kinase inhibition [27]. These hold promising avenues for targeted treatment of tumors with regulatory disruptions of PD-1 signaling.

In summary, by integrating network-based tumor stratification with pathway-centric regulatory analyses, this study provides a comprehensive view of transcriptional regulatory heterogeneity in cancer. Our findings highlight the value of gene regulatory network-based biomarkers for uncovering hidden layers of tumor biology, refining patient stratification, and guiding future research in immunotherapy and precision oncology.

## Reproducibility, data and code availability

Since the gene regulatory networks produced in this study are very large (approximately 700 GB), they have not been uploaded in full. However, all intermediate files required to reproduce the analysis are available on Zenodo at https://zenodo.org/records/17235372. The code used to reproduce the analysis and generate the figures presented in this manuscript is available on GitHub at https://github.com/kuijjerlab/PANGOLIN.

## Acknowledgments

The authors gratefully acknowledge Ieva Rauluseviciute and Romana Pop for their valuable help with the Snakemake pipeline.

## SUPPLEMENTARY FIGURES AND TABLES

**Table S1:**
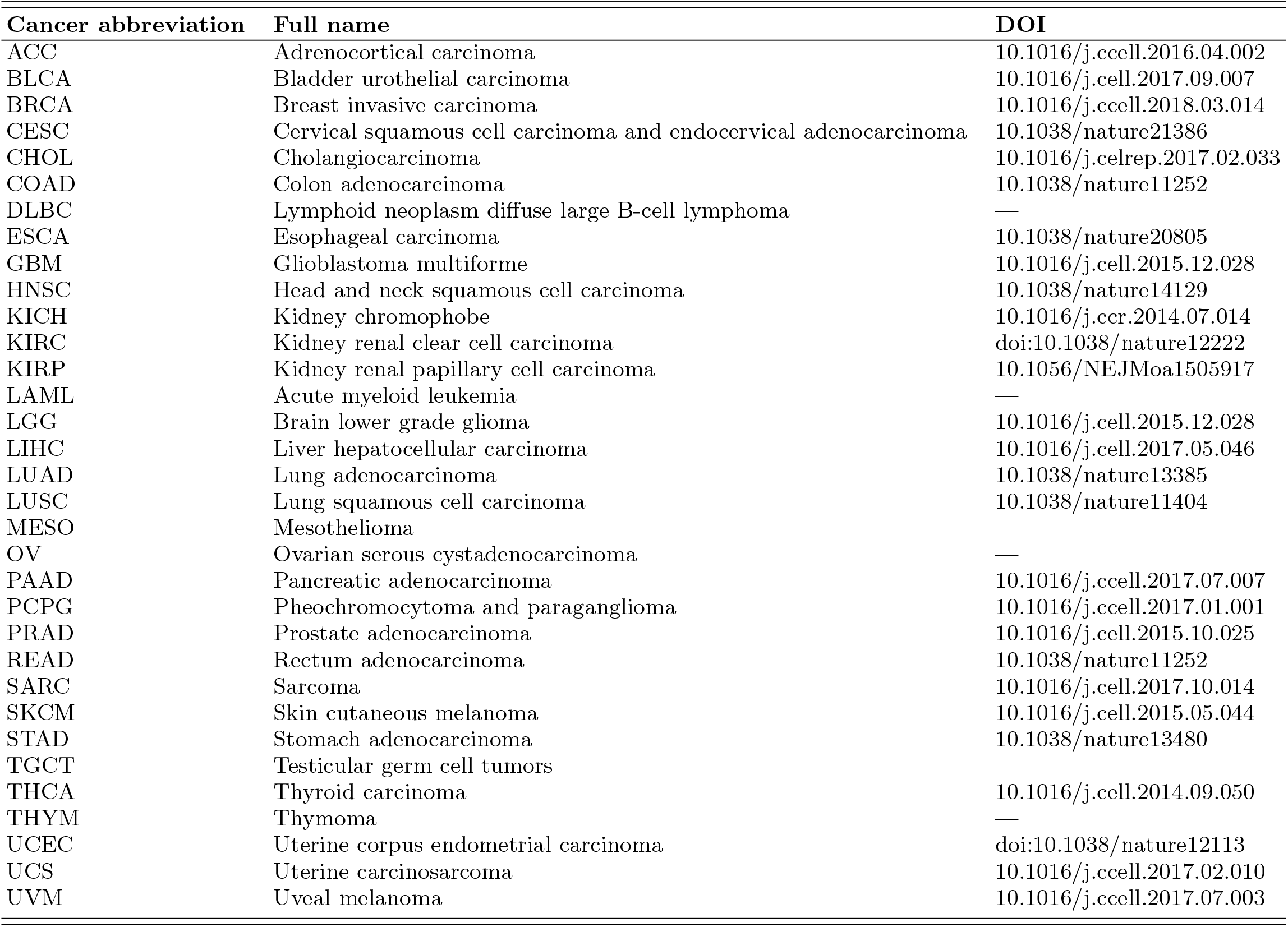
List of cancer abbreviations, their full names, and corresponding DOIs.

| **Cancer abbreviation** | **Full name** | **DOI** |
| --- | --- | --- |
| ACC | Adrenocortical carcinoma | 10.1016/j.ccell.2016.04.002 |
| BLCA | Bladder urothelial carcinoma | 10.1016/j.cell.2017.09.007 |
| BRCA | Breast invasive carcinoma | 10.1016/j.ccell.2018.03.014 |
| CESC | Cervical squamous cell carcinoma and endocervical adenocarcinoma | 10.1038/nature21386 |
| CHOL | Cholangiocarcinoma | 10.1016/j.celrep.2017.02.033 |
| COAD | Colon adenocarcinoma | 10.1038/nature11252 |
| DLBC | Lymphoid neoplasm diffuse large B-cell lymphoma | — |
| ESCA | Esophageal carcinoma | 10.1038/nature20805 |
| GBM | Glioblastoma multiforme | 10.1016/j.cell.2015.12.028 |
| HNSC | Head and neck squamous cell carcinoma | 10.1038/nature14129 |
| KICH | Kidney chromophobe | 10.1016/j.ccr.2014.07.014 |
| KIRC | Kidney renal clear cell carcinoma | doi:10.1038/nature12222 |
| KIRP | Kidney renal papillary cell carcinoma | 10.1056/NEJMoa1505917 |
| LAML | Acute myeloid leukemia | — |
| LGG | Brain lower grade glioma | 10.1016/j.cell.2015.12.028 |
| LIHC | Liver hepatocellular carcinoma | 10.1016/j.cell.2017.05.046 |
| LUAD | Lung adenocarcinoma | 10.1038/nature13385 |
| LUSC | Lung squamous cell carcinoma | 10.1038/nature11404 |
| MESO | Mesothelioma | — |
| OV | Ovarian serous cystadenocarcinoma | — |
| PAAD | Pancreatic adenocarcinoma | 10.1016/j.ccell.2017.07.007 |
| PCPG | Pheochromocytoma and paraganglioma | 10.1016/j.ccell.2017.01.001 |
| PRAD | Prostate adenocarcinoma | 10.1016/j.cell.2015.10.025 |
| READ | Rectum adenocarcinoma | 10.1038/nature11252 |
| SARC | Sarcoma | 10.1016/j.cell.2017.10.014 |
| SKCM | Skin cutaneous melanoma | 10.1016/j.cell.2015.05.044 |
| STAD | Stomach adenocarcinoma | 10.1038/nature13480 |
| TGCT | Testicular germ cell tumors | — |
| THCA | Thyroid carcinoma | 10.1016/j.cell.2014.09.050 |
| THYM | Thymoma | — |
| UCEC | Uterine corpus endometrial carcinoma | doi:10.1038/nature12113 |
| UCS | Uterine carcinosarcoma | 10.1016/j.ccell.2017.02.010 |
| UVM | Uveal melanoma | 10.1016/j.ccell.2017.07.003 |

**Supplementary Figure S1:**
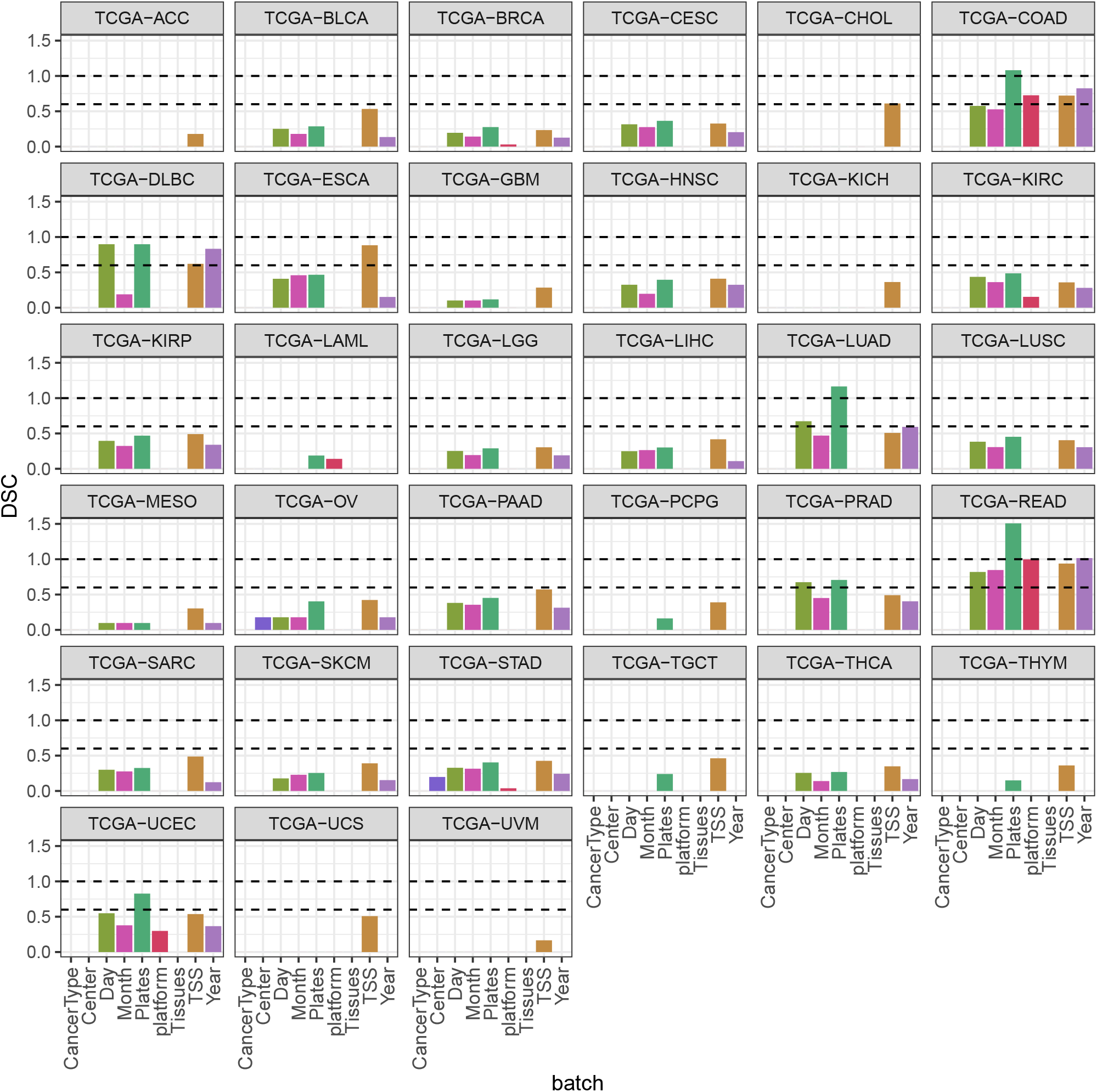
Assessment of batch effects across TCGA cancer types. Distance similarity coefficient (DSC) values were computed for each cancer type, with higher DSC values indicating stronger batch effect. Cancer types with DSC above 0.6 (lower dashed line) were further examined for potential batch effects.

**Supplementary Figure S2:**
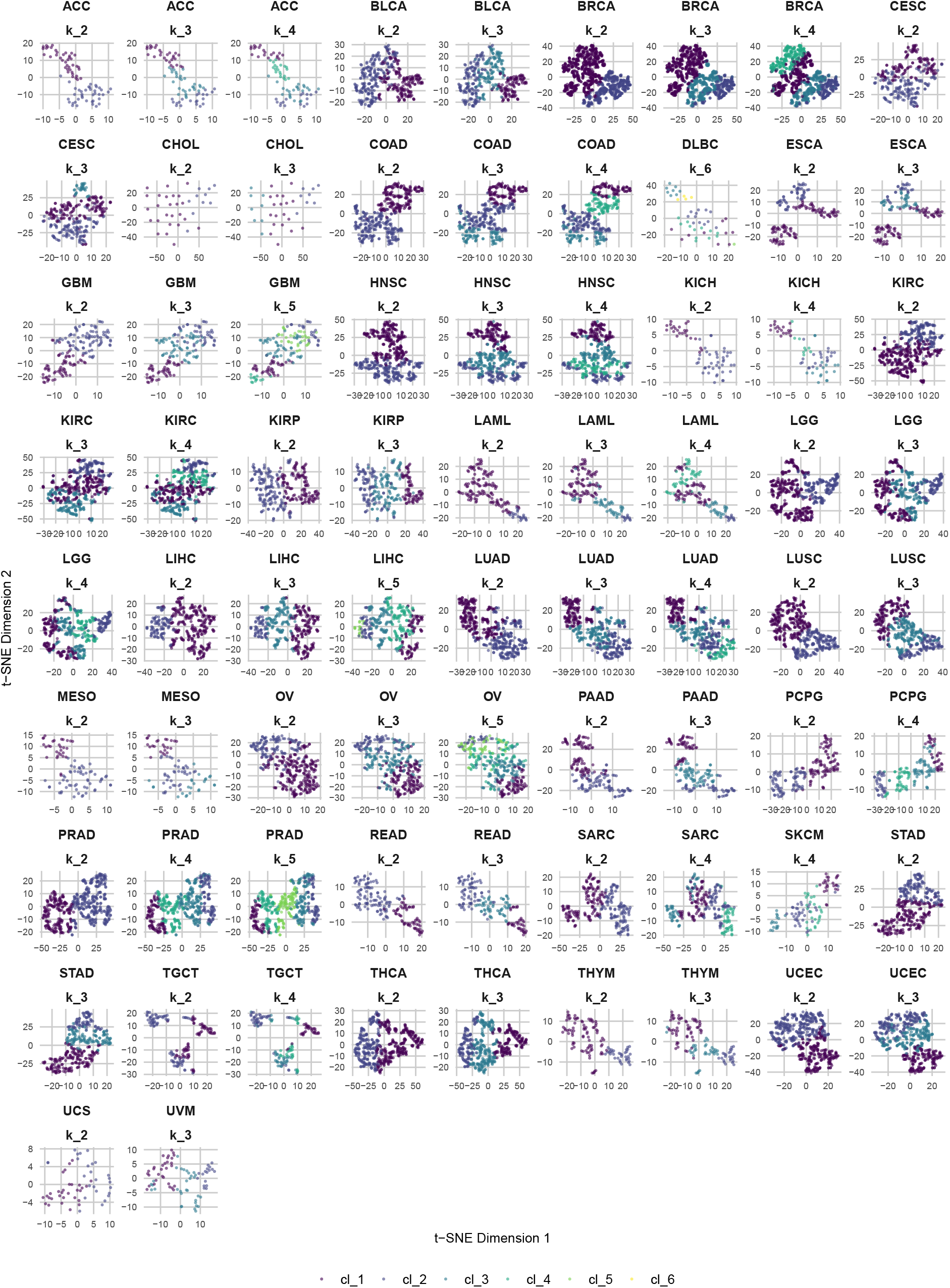
t-SNE Visualization of network-based clusters by cancer type and k (number of clusters) obtained using the unsupervised “cola” clustering algorithm.

**Supplementary Figure S3:**
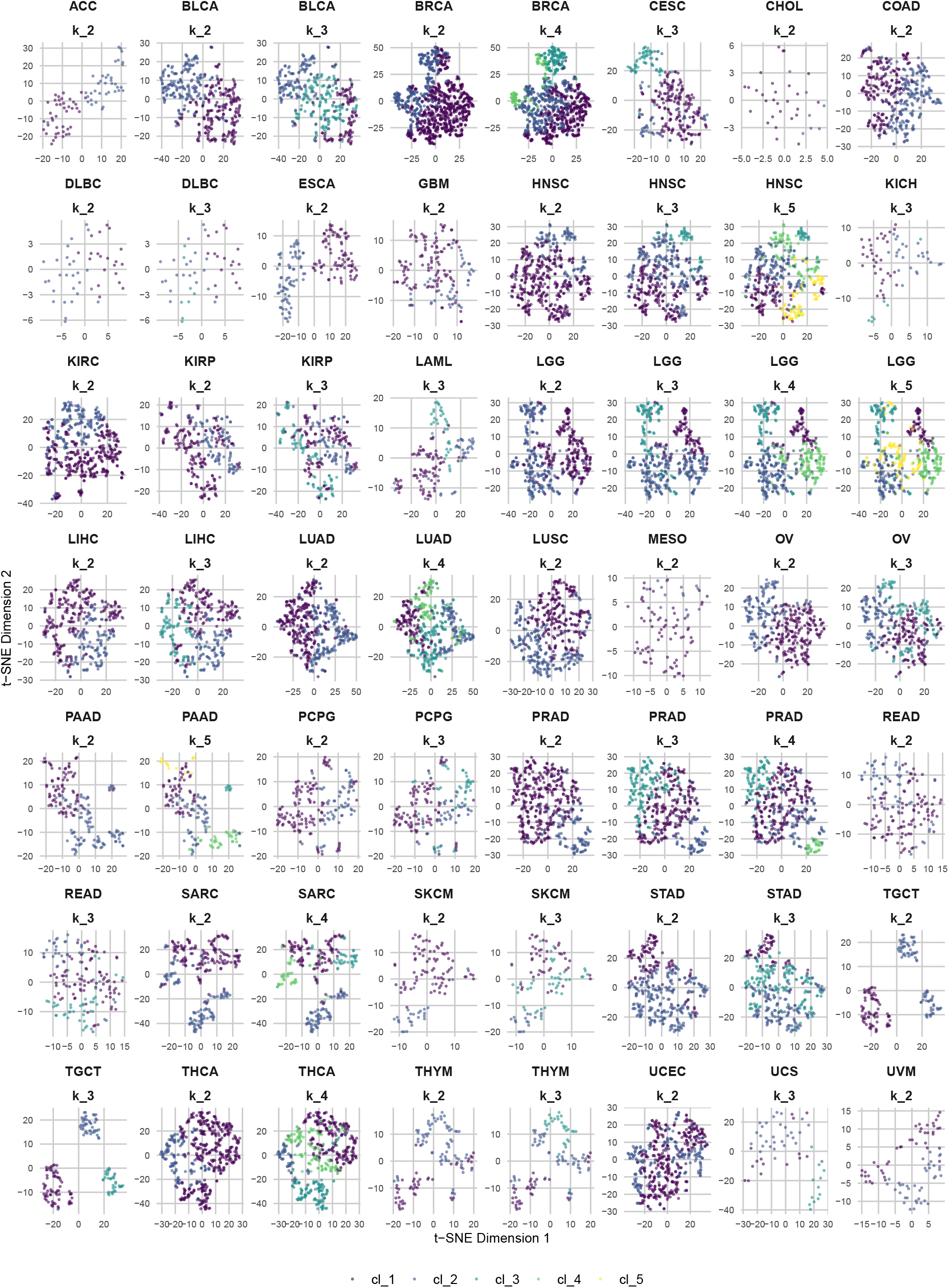
t-SNE Visualization of expression clusters by cancer type and k (number of clusters) obtained using the unsupervised “cola” clustering algorithm.

**Supplementary Figure S4:**
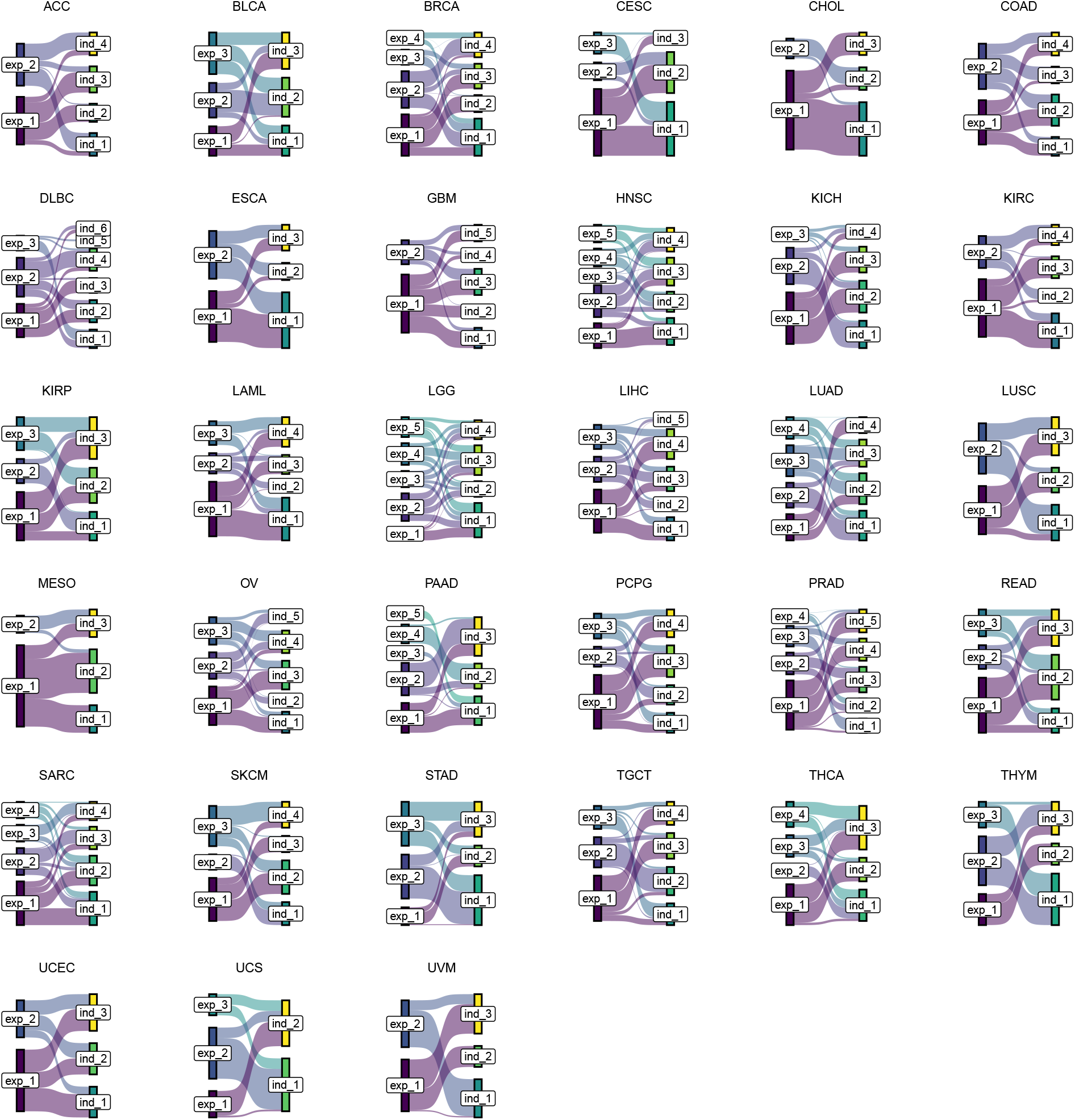
Sankey plot illustrating the comparison between expression-based clusters and network-based clusters across different cancer types.

**Supplementary Figure S5:**
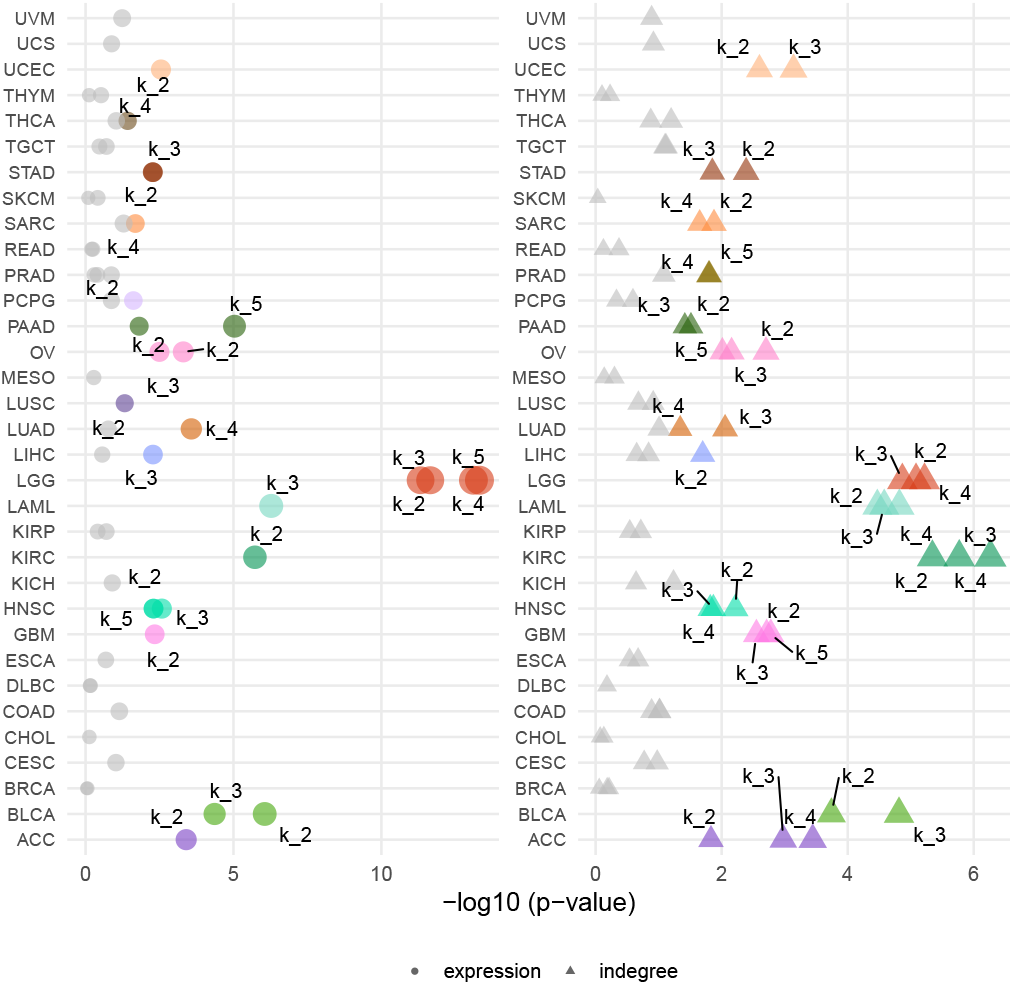
Univariate Cox regression p-values by cancer type and cluster partitions identified through “cola” clustering for expression (circle) and indegree (triangle) clusters. Significant p-values (p-value *<*0.05) are shown in color, with colors corresponding those used for the individual cancer types in Figure 2, while non-significant p-values are shown in gray. Circle and triangle sizes represent –log(p-value).

**Supplementary Figure S6:**
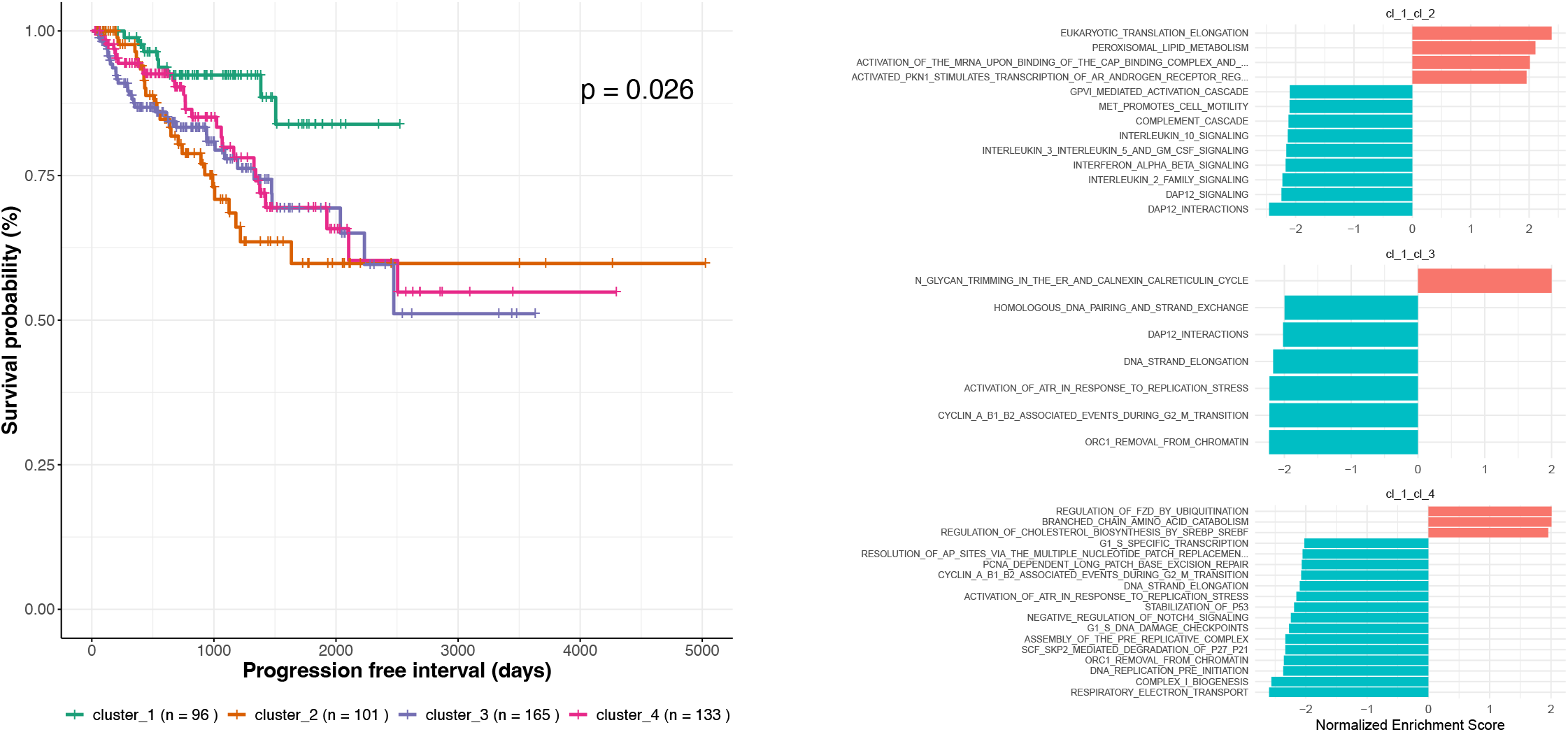
(a) Kaplan-Meier curves for the progression free survival across “cola” network-based clusters for Prostate Adenocarcinoma (PRAD) and (b) Barplot representation of the pathways that were significantly differentially regulated (FDR *<*0.05, absolute ES *>*0.5) between the cluster 1 and each of the remaining clusters.

**Supplementary Figure S7:**
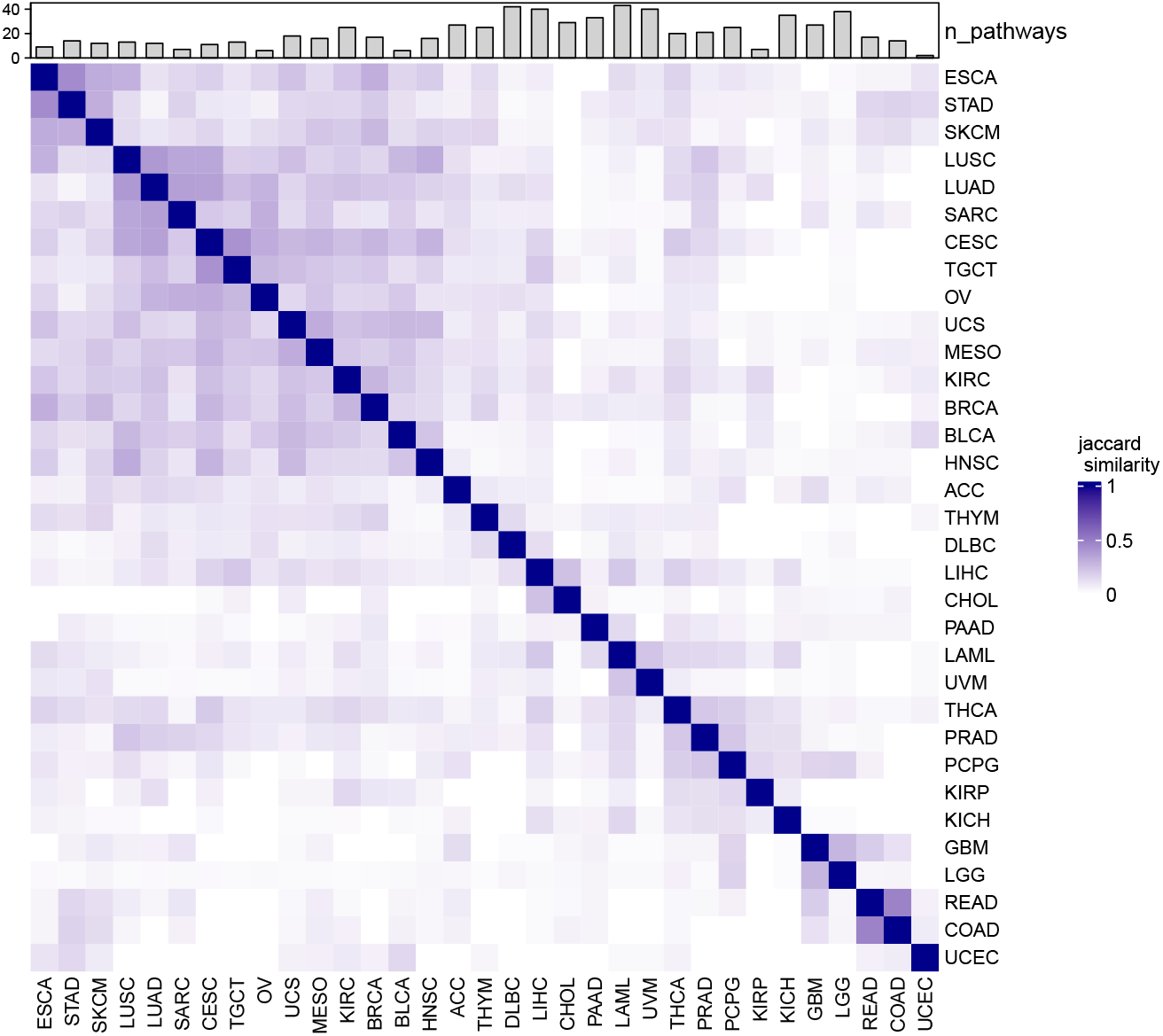
Jaccard similarity between the lists of significant pathways (selected based on an FDR *<* 0.05, variance explained by the first principal component above 25% and effect sizes above the 95th percentile) for each pair of cancer types.

**Supplementary Figure S8:**
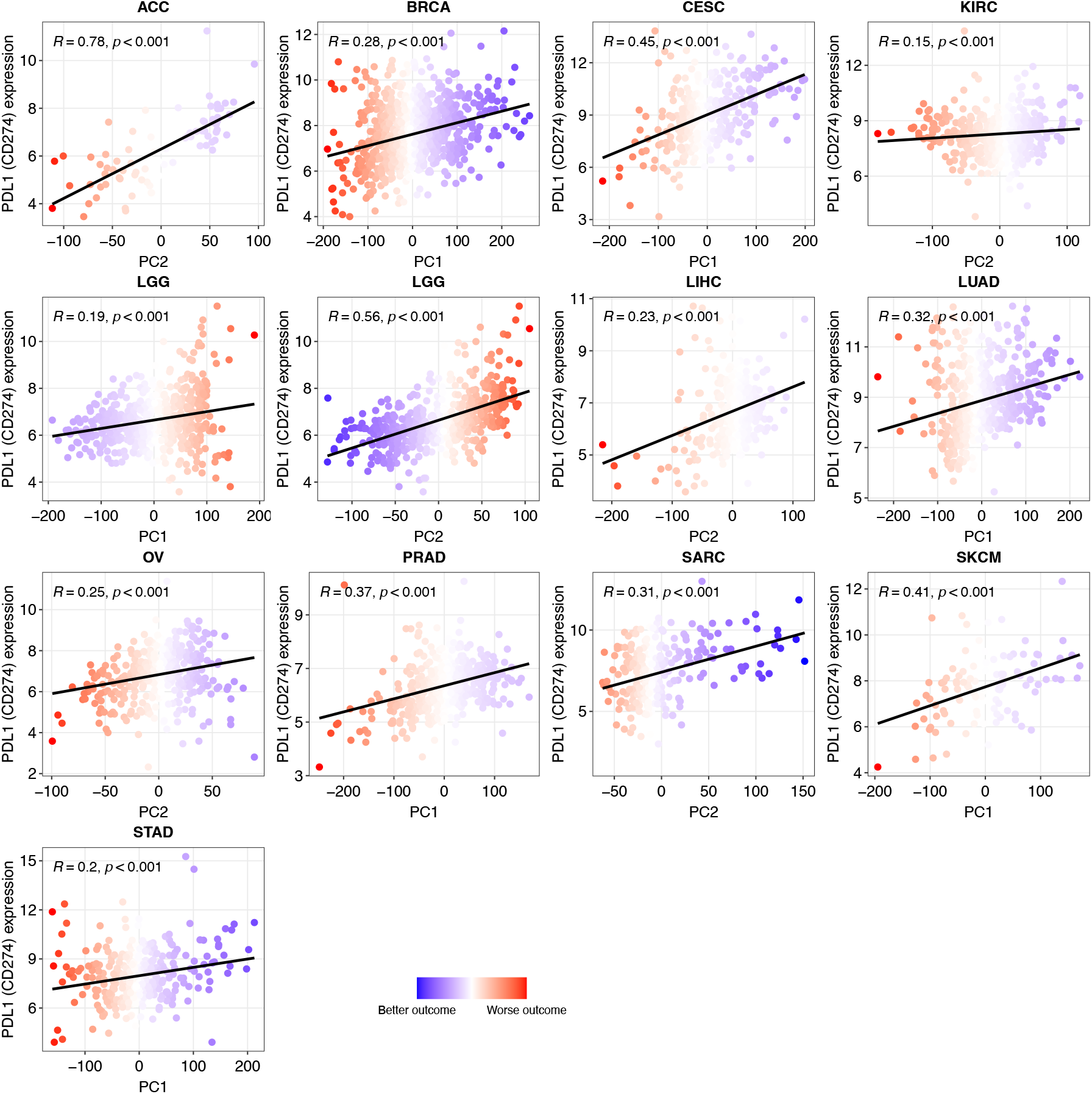
Spearman correlations between PD-L1 (*CD274*) expression levels and the principal component significantly associated with survival outcomes. Each dot is colored based on the predicted hazard ratio derived from the univariate Cox regression model.

**Supplementary Figure S9:**
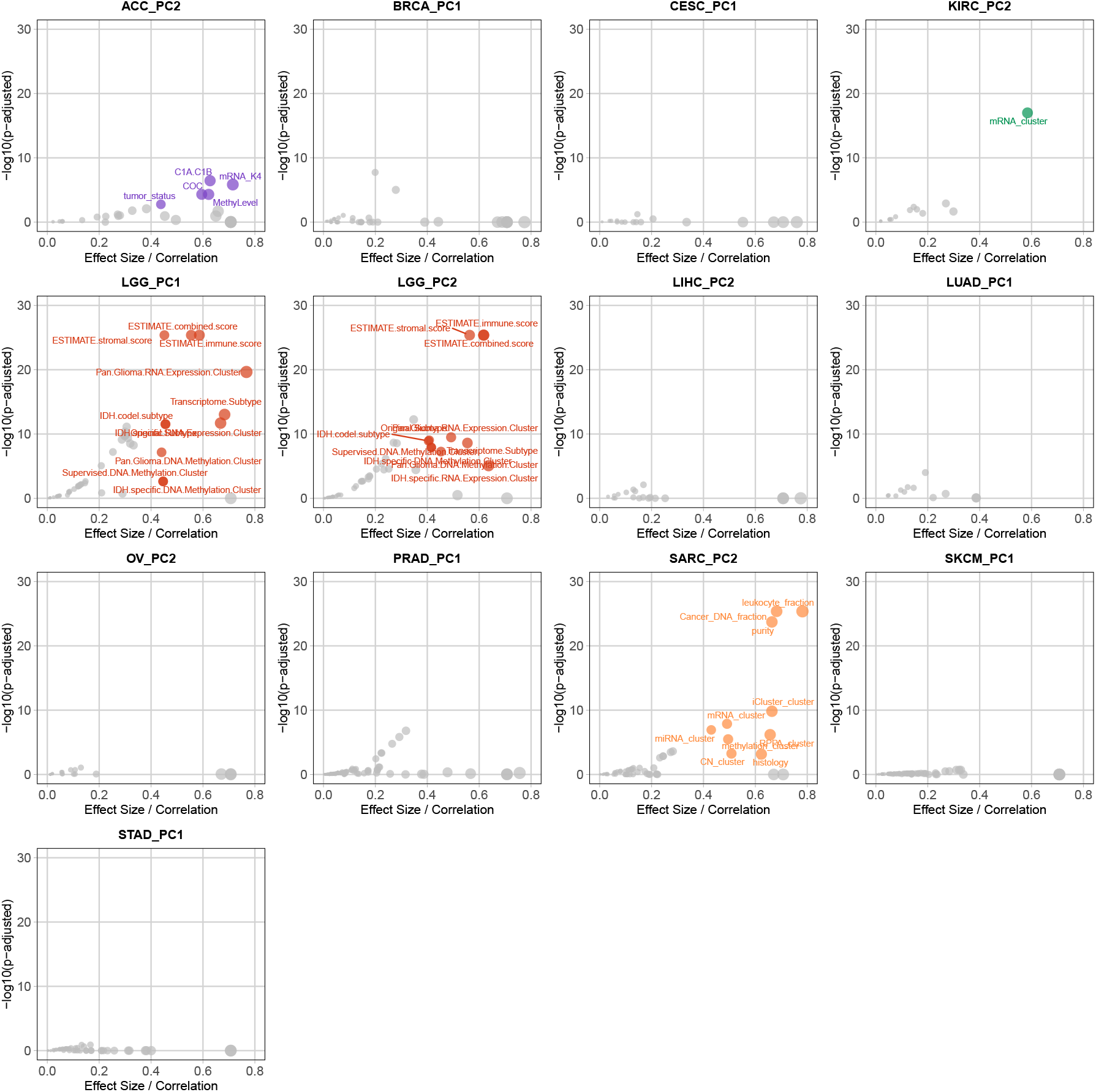
Associations between the PD-1 pathway-based patient heterogeneity scores (based on PORCUPINE PC1, PC2, or both, depending on the cancer type) and clinical features and “omics” subtypes in selected cancer types. Significant associations (rank-biserial correlation or Spearman correlation above 0.4 and *FDR <* 0.01) are indicated with colors using the same color coding as in Figure 2, while non-significant associations are shown in gray.

**Supplementary Figure S10:**
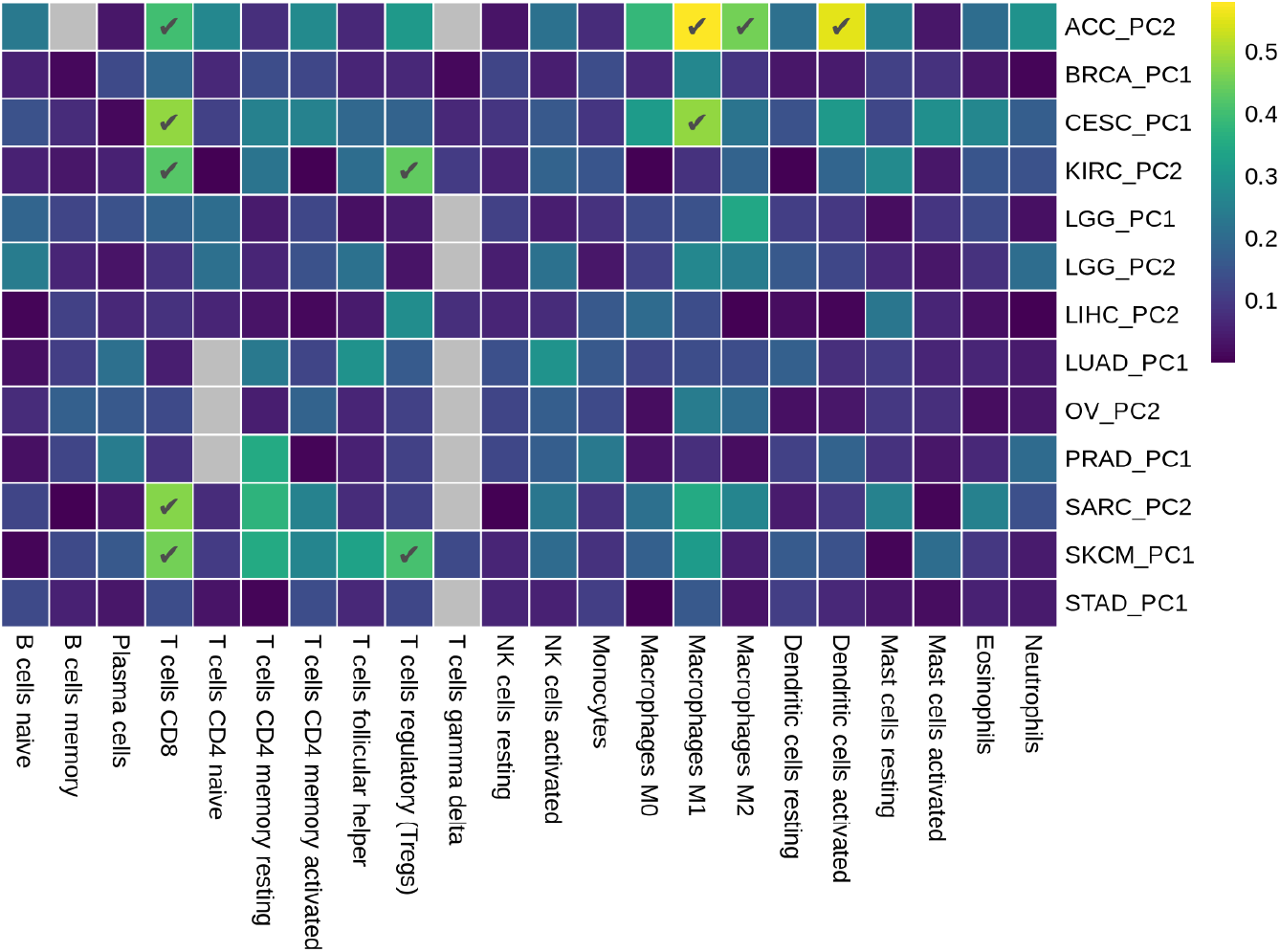
Spearman correlations (color bar) between those PD-1 pathway-based patient heterogeneity scores that associate with survival (based on PORCUPINE PC1, PC2, or both, depending on the cancer type) and the estimated proportions of immune cells in the twelve cancer types where pathway heterogeneity scores significantly associated with survival. Correlations above 0.4 are marked by checkmarks (✓).

**Supplementary Figure S11:**
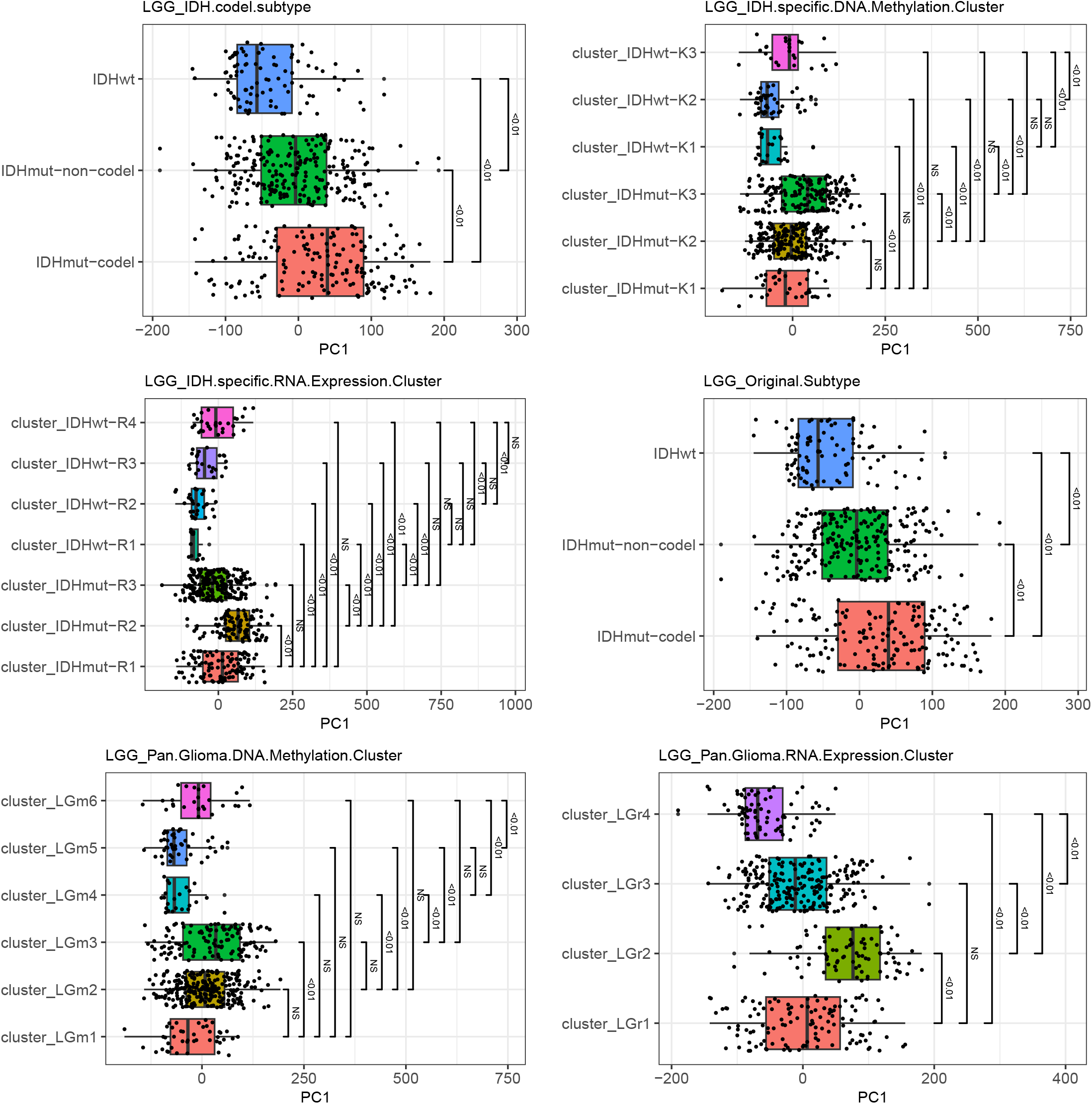
Comparison of PD-1 pathway-based patient heterogeneity scores across clinical and molecular subtypes in brain lower grade glioma (LGG) where PD-1 regulatory heterogeneity (based on PORCUPINE PC1) is significantly associated with clinical outcome. Annotations correspond to those provided in [49]. False discovery rates from pairwise group comparisons are displayed; NS non-significant.

**Supplementary Figure S12:**
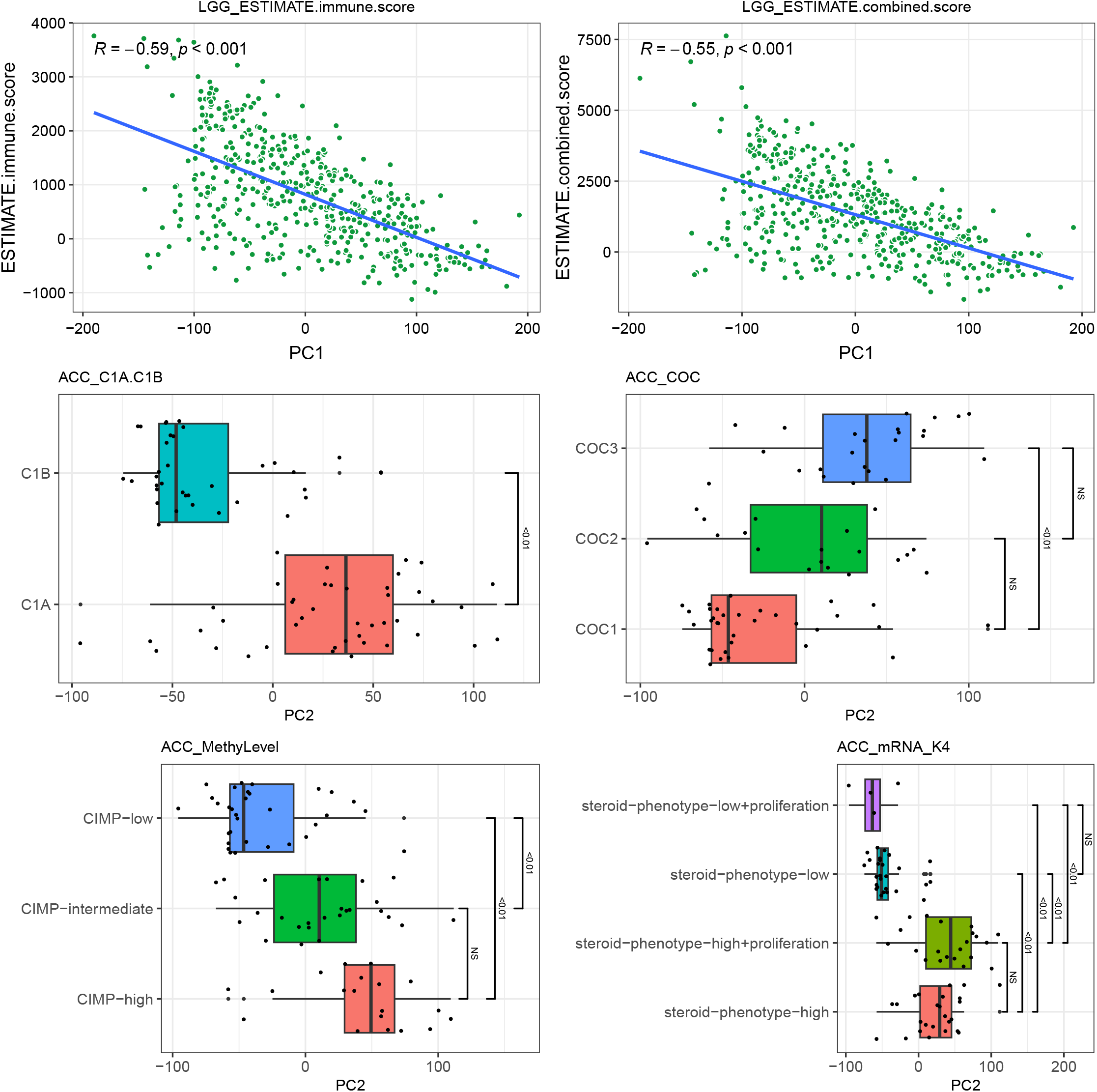
Comparison of PD-1 pathway-based patient heterogeneity scores across clinical and molecular subtypes in brain lower grade glioma (LGG) and adrenocortical carcinoma (ACC) where PD-1 regulatory heterogeneity (based on PORCUPINE PC1 in LGG and PC2 in ACC) is significantly associated with clinical outcomes. Annotations correspond to those provided in [49] and in [50]. False discovery rates from pairwise group comparisons are displayed; NS non-significant

**Supplementary Figure S13:**
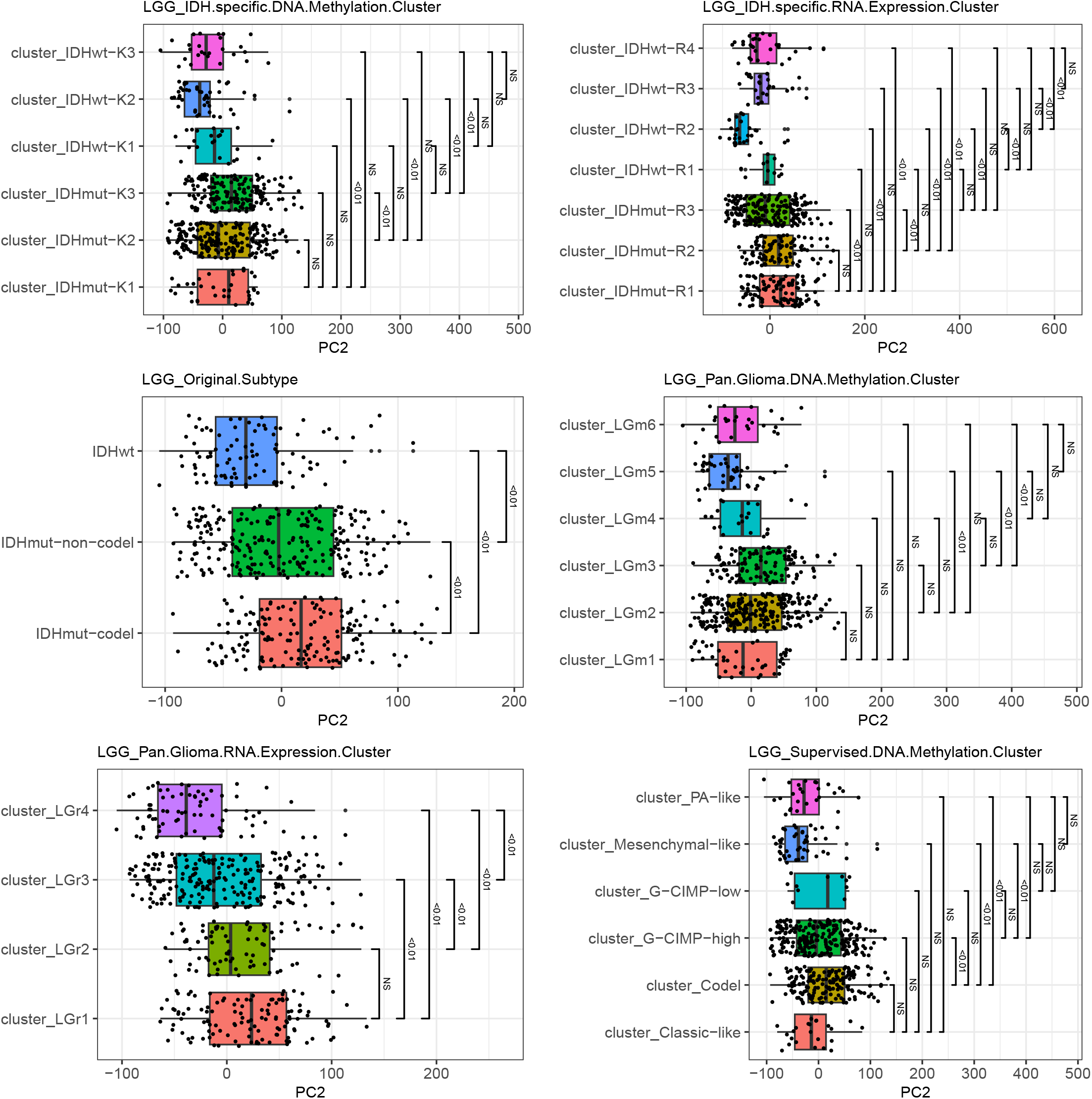
Comparison of PD-1 pathway-based patient heterogeneity scores across clinical and molecular subtypes in brain lower grade glioma (LGG) where PD-1 regulatory heterogeneity (based on PORCUPINE PC2) is significantly associated with clinical outcome. Annotations correspond to those provided in [49]. False discovery rates from pairwise group comparisons are displayed; NS non-significant.

**Supplementary Figure S14:**
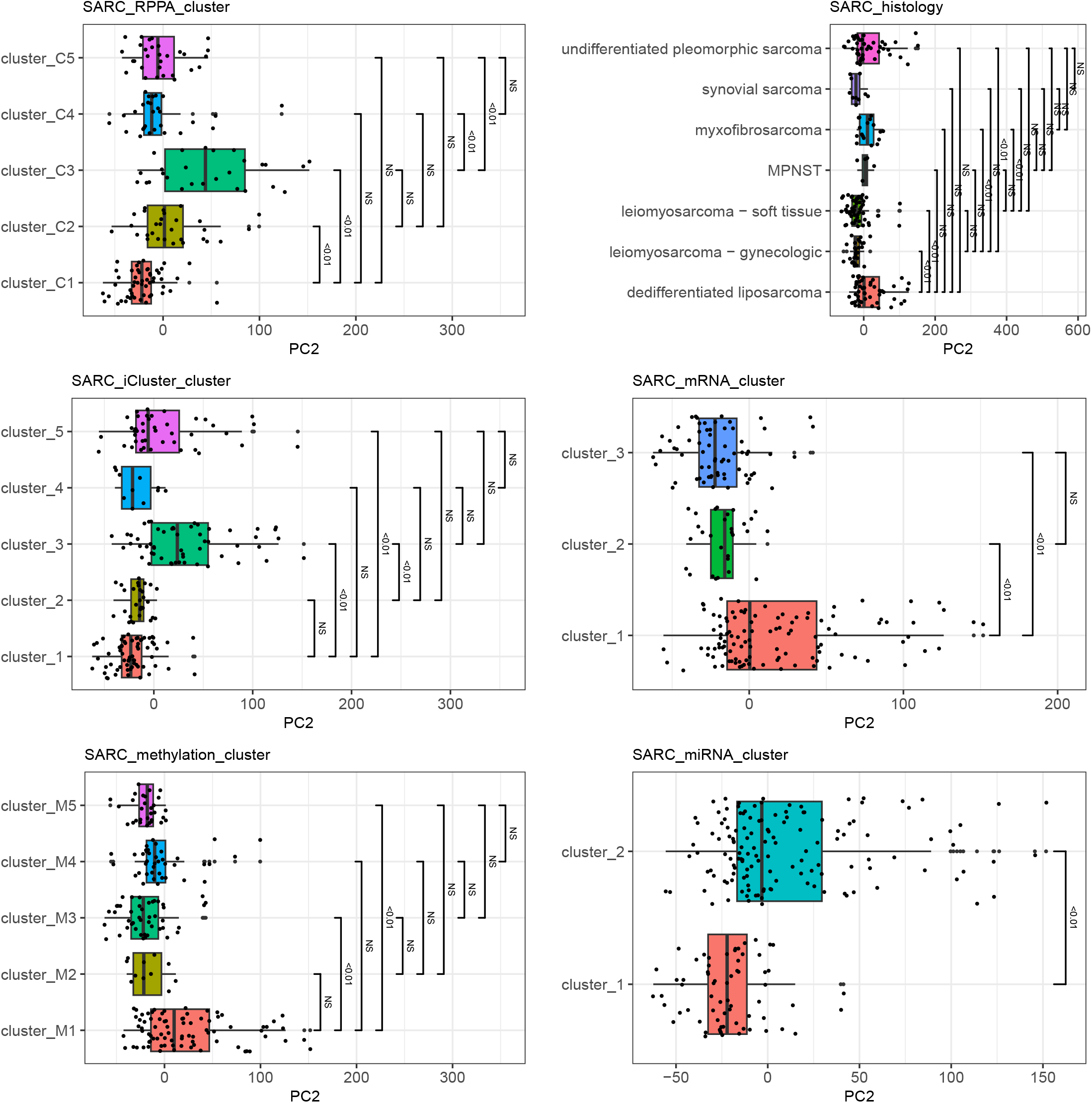
Comparison of PD-1 pathway-based patient heterogeneity scores across clinical and molecular subtypes in sarcoma (SARC) where PD-1 regulatory heterogeneity (based on PORCUPINE PC2) is significantly associated with clinical outcome. Annotations correspond to those provided in [51]. False discovery rates from pairwise group comparisons are displayed; NS non-significant.

**Supplementary Figure S15:**
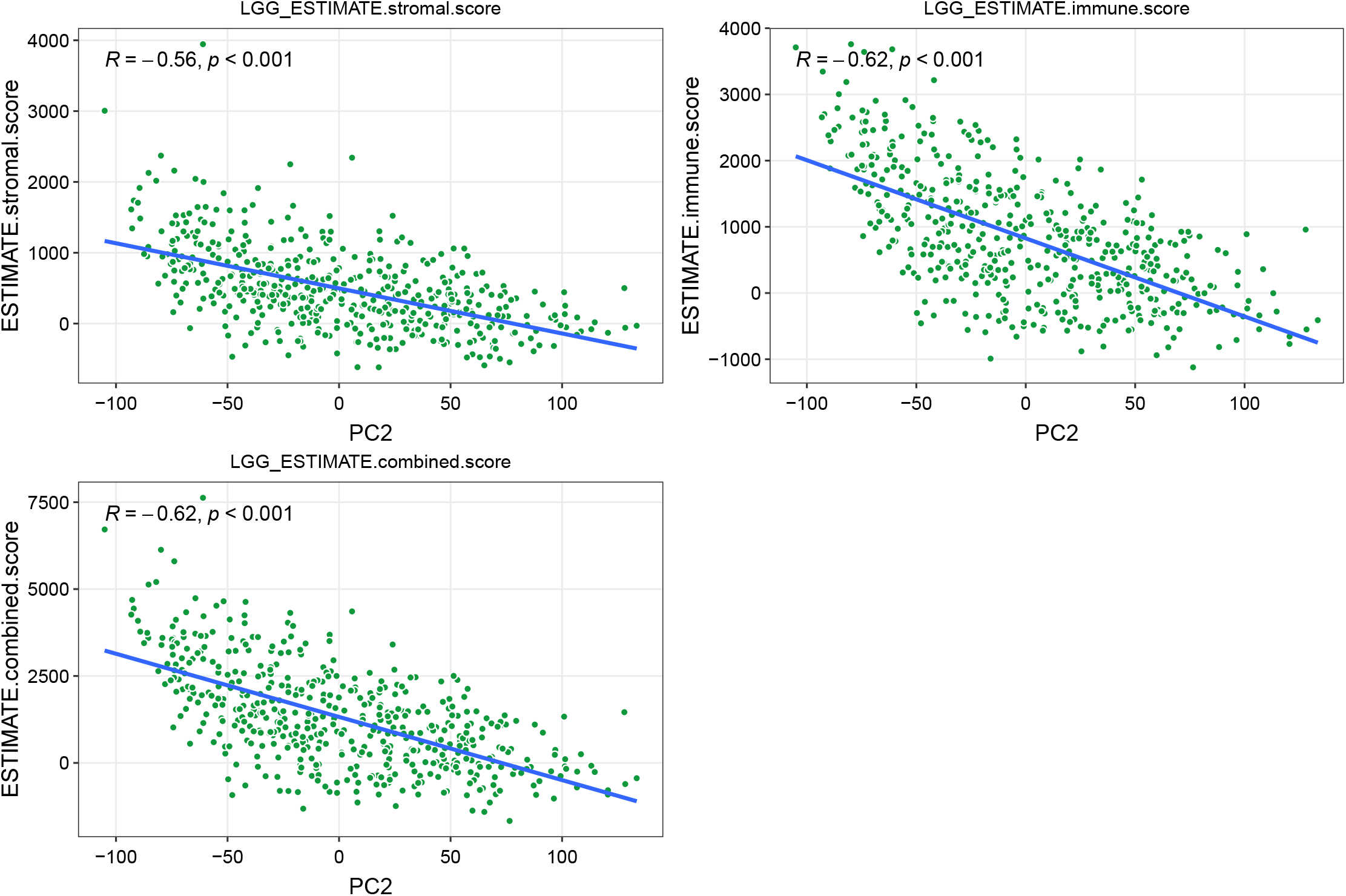
Comparison of PD-1 pathway-based patient heterogeneity scores across clinical and molecular subtypes in brain lower grade glioma (LGG) where PD-1 regulatory heterogeneity (based on PORCUPINE PC2) is significantly associated with clinical outcome. Annotations indicate clusters as defined in [49]. False discovery rates from pairwise group comparisons are displayed; NS non-significant.

**Supplementary Figure S16:**
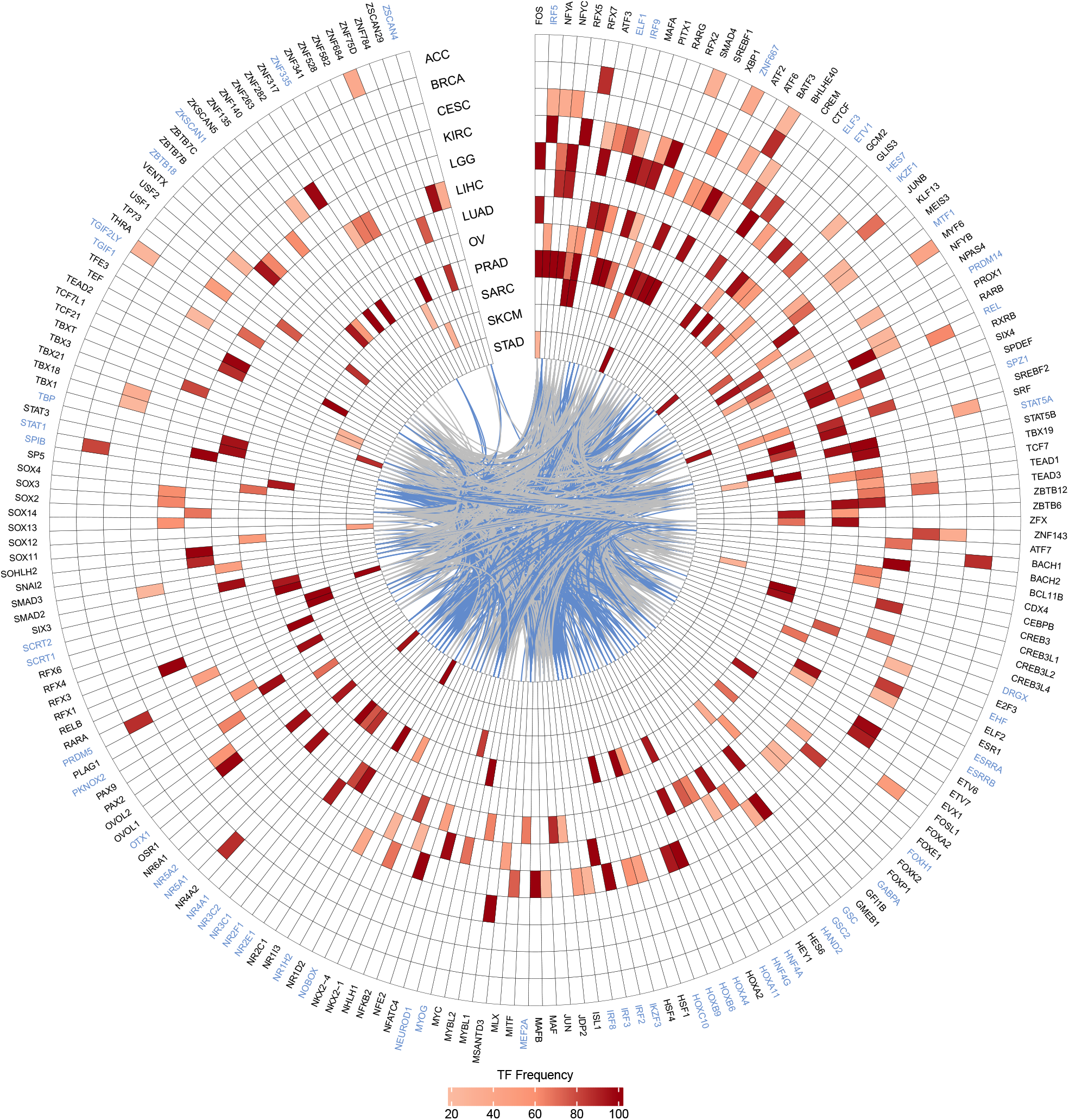
TFs selected through the application of the regularized Cox regression model to edge weights between 671 TFs and the *CD274* gene across various cancer types. TFs with non-zero coefficients were selected and were predictive in at least 20 out of 100 model runs (compared to 50 out of 100 model runs in Figure 4). The color of each cell reflects the frequency with which the TF was selected as predictive. Grey inner connections represent known interactions among TFs based on protein-protein interaction (PPI) data, while blue connections indicate TFs predicted to regulate *CD274* according to the motif prior network. Blue TF names indicate these TFs have prior binding sites in the promoter of *CD274*.

